# Remodeling of purinergic signaling in the paraventricular hypothalamus promotes hyperphagic obesity and insulin resistance

**DOI:** 10.1101/2024.07.02.601503

**Authors:** Yu Liu, Anastasiia Bondarenko, Haiyang Zhang, Pengwei Luan, Qian Zhang, Bo Wang, Syeda Sadia Najam, Xian Liang, Ali Alamdar Shah Syed, Dongfa Lin, Shangyuan Huang, Ruslan Konovalov, Witold Konopka, Zhi Zhang, Yi Zhang, Shihao Wu, Miao Jing, Ilya A. Vinnikov

## Abstract

Body energy homeostasis is tightly regulated by hypothalamic neural circuits. However, their remodeling upon metabolic challenges remains incompletely characterized, thus complicating the development of safe medications against the surge of metabolic diseases. Oxytocin (OXT) neurons in the paraventricular nucleus of the hypothalamus (PVH) are one of the key effectors regulating energy balance. In this work, we report that high-fat diet (HFD) feeding in mice evokes spatiotemporally selective ATP release from astrocytes (Inflares) in the PVH, accompanied by the expression of hematopoietic lineage-specific ADP/ATP receptor P2Y_12_ on local OXT (PVH^OXT^) neurons. Strikingly, increased purinergic signaling leads to impaired responsiveness of PVH^OXT^ neurons accompanied by hyperphagic obesity and insulin resistance in mice. Conversely, loss of P2Y_12_ in PVH^OXT^ neurons attenuates metabolic phenotypes, illustrating that such remodeling is both necessary and sufficient to induce metabolic phenotypes. Inflares were also induced by hyperglycemia, while emergence of P2Y_12_ on OXT neurons of patients with diabetes mellitus suggests an evolutionary conserved purinergic response to various metabolic challenges and its potential as a drug target. Accordingly, nasal administration of clinically approved doses of P2Y_12_ inhibitors counteracts obesity in mice and non-human primates, paving the way for application of these compounds in patients with metabolic disorders.

## INTRODUCTION

More than one billion people suffer from obesity ^1^, which contributes to associated metabolic disorders, such as diabetes mellitus. Pharmacological approaches to treat these conditions include inhibitors of pancreatic lipases, agonists of receptors for glucagon-like peptide-1 (GLP-1) and for glucose-dependent insulinotropic polypeptide (GIP) ^2,3^. However, the requirement of long-term application of these compounds for weight maintenance is associated with potential side effects ^4^, underscoring a pressing need for identifying novel targets and more effective drug delivery strategies ^5^.

Within the brain, the key neuronal circuits regulating energy homeostasis are located in the hypothalamus ^6,7^. A metabolically critical subset of second order neurons in the paraventricular nucleus of the hypothalamus (PVH) expressing MC4R ^8–13^ or other GPCR receptors senses and integrates metabolic-related signaling to release oxytocin (OXT), thus suppressing appetite and maintaining energy homeostasis ^14–18^. One of the key questions in the field is to delineate how diverse signals modulate the activity of PVH^OXT^ neurons thus enabling the dynamic adaptations upon metabolic challenges.

Metabolic disorders have been found to be associated with the dysfunction in purinergic system ^19^, which is composed of versatile extracellular ligands and membrane receptors mediating cell-cell communication ^20–22^. Recently, we identified a stress-evoked spatiotemporally selective pattern of ATP release from astrocytes, named Inflares, representing an active signal transmission system that encodes brain injury properties ^23^. The matching between injury severity and astrocyte-released Inflare numbers ensures the adequate damage management, emphasizing the importance of purinergic signaling, as well as its alteration, which might contribute to disease progression ^23,24^. Such purinergic signaling system might also exist in the hypothalamus, reacting on metabolic challenges and encoding stress information to affect the activity of ATP-sensing cells. At the receiver side, a major ATP/ADP receptor P2Y_12_, encoded by the *P2ry12* gene, was found to be exclusively restricted to microglia ^25–29^. A direct binding and activation of the *P2ry12* gene promoter by its upstream transcriptional regulator NF-κB is known to be stimulated by metabolic challenges ^30^, which also activate this transcription factor in microglia ^31,32^, hypothalamic neurons ^33,34^ and other cells. For example, NF-κB is inhibited by glucocorticoid receptor (GR) ^35^, which is highly abundant within hypothalamic neurons and is critical for energy homeostasis ^36^. Until now, the ectopic expression of neuronal P2Y_12_ in relevant hypothalamic circuits and its contribution in decoding extracellular purinergic signaling in response to metabolic challenges remained uncharacterized.

Given the tight link between purinergic signaling and metabolic diseases ^19,22^, while keeping in mind the need of identifying novel therapeutic strategies for their treatment, this work aimed to delineate the pathophysiological mechanisms, metabolic function, and relevance of the purinergic pathway within the hypothalamus.

## RESULTS

### Metabolic challenges stimulate astrocytic ATP release within the PVH

First, we tested whether, in analogy with other types of brain injuries ^23^, Western diet or diabetes may induce ATP Inflares within the PVH, the critical region regulating energy intake and expenditure. We stereotaxically injected recombinant adeno-associated viral vector (rAAV) encoding the ATP1.0 sensor under astrocyte-specific promoter into the PVH of adult mice fed with high-fat diet (HFD) or normal chow food (**Figure 1A**). Next, using two-photon microscopy, we recorded the extracellular ATP dynamics in acute PVH slices from these mice. Remarkably, starting from as early as two weeks of HFD feeding, we observed a progressive increase in the frequency of ATP events (Inflares) in the PVH compared to unchanged levels in chow diet-fed animals (**Figure 1B,C** and **Movies S1-S2**). Unaltered amplitude, size and duration of individual Inflares between time points of HFD feeding are consistent with our recent report that individual Inflares serve as the fundamental signaling units (**Figure S1A-C**). Notably, pre-treatment with probenecid, the inhibitor of the pannexin 1 channel, reduced the Inflare frequency in HFD-treated mice (**Figure 1A,D,E** and **Movie S3**), confirming the hemichannel-dependent ATP release ^23^. To further study the mechanism and origin of the ATP release, we selectively knocked-out pannexin 1 in astrocytes by using GFAP:Panx1cko mice. This led to a significant decrease of the frequency, but not amplitude, size or duration of Inflares compared to the contralateral PVH stereotaxically injected with the control rAAV equipped with a construct without Cre recombinase (**Figures 1F-H**, **S1D-F** and **Movie S4**), supporting astrocytes as the emitters of ATP release and its dependence on pannexin 1, consistent with the mechanism of injury-evoked Inflares ^23^. Due to the identical molecular mechanism, laser injuries in the PVH slices from HFD-treated mice failed to cause additional elevation of population frequency or changes in individual properties of Inflares (**Figures 1H, S1D-J**).

**Figure 1.**
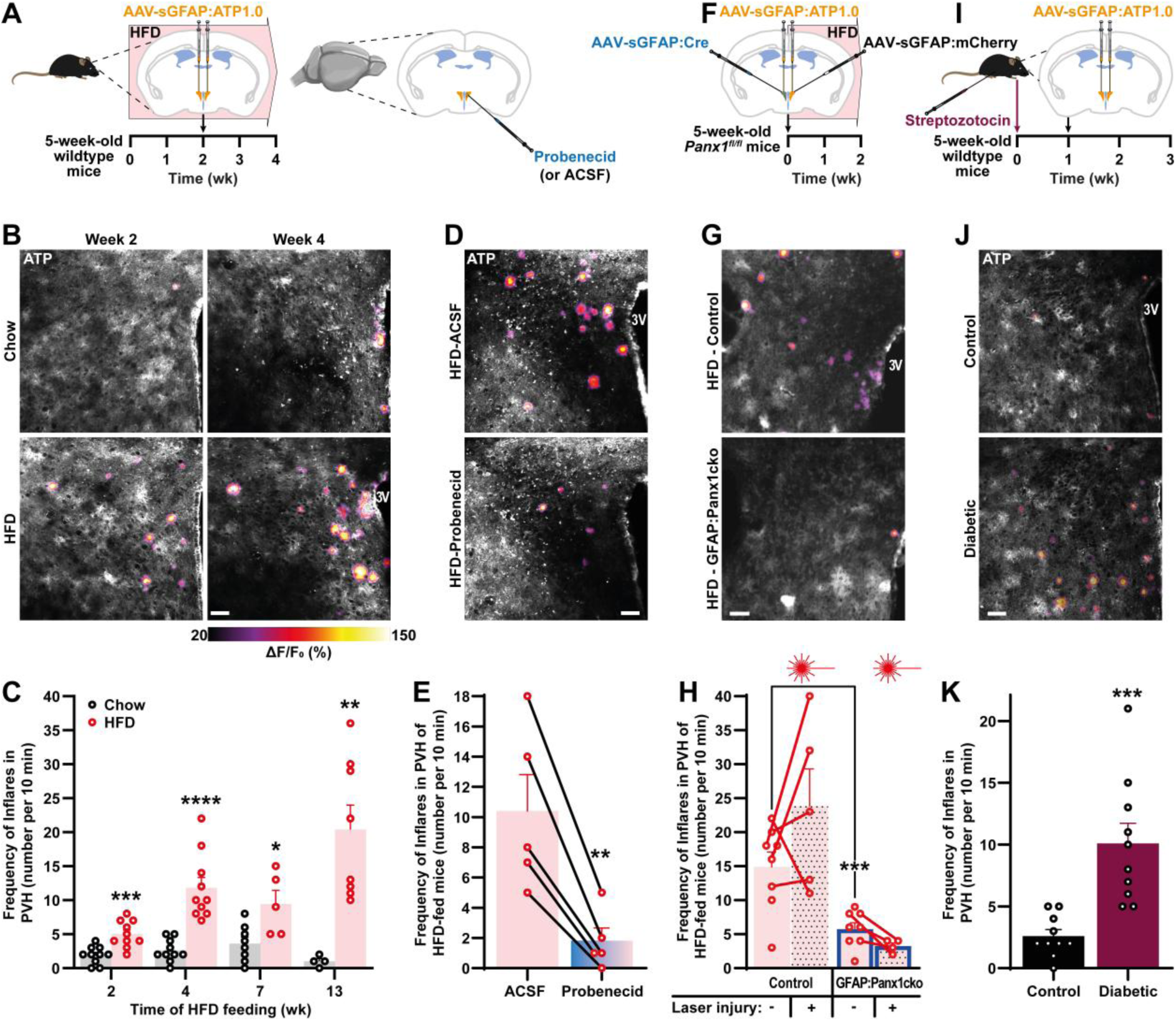
HFD or hyperglycemia induce ATP release from astrocytes in PVH. (**A-C**) Experimental design (**A**), representative microphotographs (**B**) and quantification (**C**) of the ATP Inflares in the PVH regions-containing acute hypothalamic slices of male wild type mice fed with chow or high-fat diet (HFD) for indicated time periods (2 and 4 wk analyses: n = 10 from 3 mice per group; 7 wk: n = 8 and 5 from 3 mice; 13 wk: n= 4 and 8 from 3 mice for Chow and HFD groups, respectively). (**D-E**) Representative microphotographs (**D**) and quantification (**E**) of Inflares in the PVH-containing acute slices of male mice fed with HFD for 4 wk before and after treatment with probenecid (n = 5 slices from 3 mice). (**F-H**) Male *Panx1^fl/fl^* mice were stereotaxically injected to one PVH side with recombinant adeno-associated viral vector (rAAV) equipped with GFAP-driven Cre (GFAP:Panx1cko) and to the other side—with control rAAV followed by HFD-feeding for 2 wk. After recording of ATP events, some slices have been subjected to focal laser injury followed by repeated recording and analysis. Experimental design (**F**), representative microphotographs (**G**) and quantification (**H**) of Inflares in the PVH-containing acute slices (n = 8 slices from 3 mice, out of which 4 and 5 slices were also laser-irradiated on the Cre and control sides, respectively). (**J-K**) Experimental design (**I**), representative microphotographs (**J**) and quantification (**K**) of Inflares in male mice 2 wk after streptozotocin or vehicle injection (n = 10 slices from 3 mice from each group). The peak fluorescence change (ΔF/F_0_) of Inflares recorded by the ATP1.0 sensor over 10 minutes are pseudocolored for better visualization. The frequency of Inflares over 10 min recording were quantified. ACSF, artificial cerebro-spinal fluid. Error bars represent standard error of means (SEM). *, *p* < 0.05; **, *p* < 0.01; ***, *p* < 0.001; ****, *p* < 0.0001 as analyzed by unpaired (**C**,**K**) or paired (**E,H**) two-tailed Student’s t-tests. Scale bar in 50 µm. See also **Figure S1**.

To explore whether the up-regulation of purinergic signals represent general strategy in response to metabolic challenges, we analyzed the ATP dynamics in the PVH of a mouse model of type I diabetes mellitus induced by streptozotocin ^37^. Consistent with the HFD model, we observed an elevated frequency of Inflares in the PVH of diabetic mice (**Figure 1I-K** and **Movie S5**), confirming that metabolic stressors induce selective patterns of astrocytic ATP release that contributes to altered purinergic signaling within the PVH.

### Emergence of P2Y_12_ on PVH^OXT^ neurons upon metabolic challenges across species

Next, we sought to investigate how these Inflares signal within the PVH of metabolically compromised mice. Strongly enriched in this region ^38^ and reported to be exclusively expressed in microglia within the central nervous system ^26,39^, P2Y_12_ is a Gα_i_-protein coupled receptor that could sense ATP/ADP in Inflares. Strikingly, HFD induced an expression of this microglia-specific receptor on 54.67 ± 3.62% of PVH^OXT^ neurons (**Figure 2A**) compared to no expression in PVH^OXT^ neurons of chow-fed animals (*p* < 0.0001). To validate these results, we analyzed the level of *P2ry12* transcript in FACS-isolated PVH^OXT^ neurons from HFD-fed mice prior to the onset of obesity (**Figure S2A**) and confirmed its drastic up-regulation compared to the level in PVH^OXT^ neurons from chow-diet fed animals (**Figure 2B**). Analysis of recent single cell RNA-seq database (https://db.cngb.org/cdcp/hca/) consistently reports the emergence of *Oxt*^+^*P2ry12*^+^ double-positive neurons in mice fed with HFD (**Figure 2C**) ^40–42^. Moreover, in line with previous studies ^33,34^, HFD treatment also stimulated OXT neurons of non-human primates to express NF-κB transcriptional regulator ^43^, which is known to directly activate *P2ry12* transcription ^30^ (**Figure S2B**) implying the generalization of HFD-induced purinergic remodeling across species.

**Figure 2.**
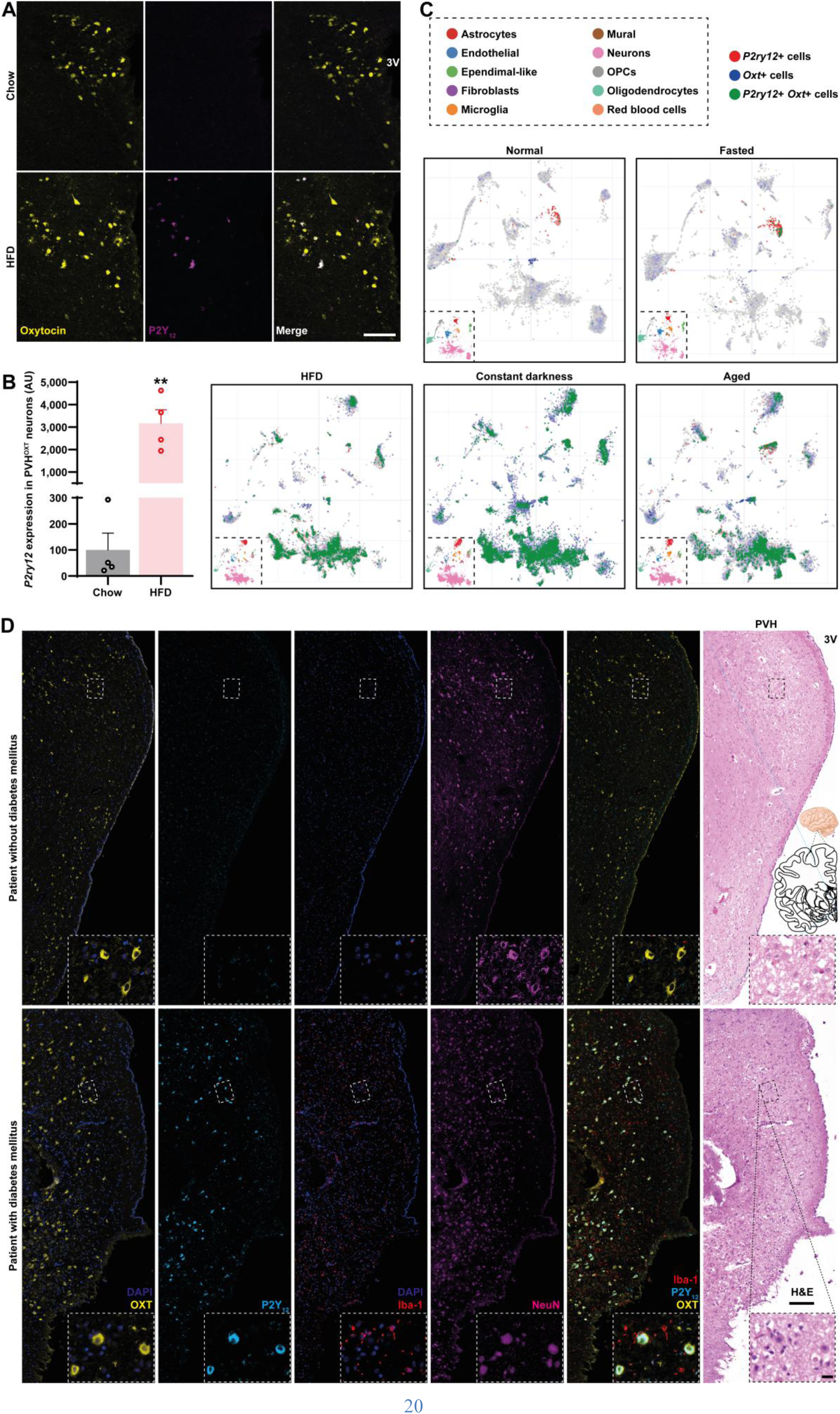
Expression of P2Y_12_ on PVH^OXT^ neurons of HFD-fed mice and patients with diabetes mellitus. (**A**) Representative confocal microphotographs of the hypothalamic slices immunostained for P2Y_12_ from OXT^IRES-Cre^-driven reporter male mice fed with HFD for 10 wk (n = 4). (**B**) qRT-PCR analysis of *P2ry12* in FACS-isolated PVH^OXT^ neurons of female mice fed with HFD for 4 wk (n = 4). (**C**) Co-localization (green dots) of *P2ry12* (red dots) with *Oxt* (blue dots) in the indicated hypothalamic cellular populations (insets) of aged (19-24 months) and young (3 months) mice subjected to 24 h-fasting, constant darkness or fed with normal or HFD food (assessed by https://db.cngb.org/cdcp/hca/). OPCs, oligodendrocyte precursor cells. Note that the cellular clusters in the cell maps (insets) are not completely homogenic, but contain many inclusions outlined with a different color, i.e. a small number of cells of different populations within the predominant ones, which, however, cluster together. (**D**) Representative co-localization microphotographs of the PVH-containing *ex vivo* slices from patients with or without diabetes mellitus (n = 3 and 2, respectively). Error bars represent standard error of means (SEM). **, *p* < 0.01 as analyzed by unpaired two-tailed Student’s t-test. Scale bar in µm: 100 (**A**), 200 (**D**) and 20 (insets). See also **Figure S2**.

Additionally, we examined whether similar emergence of P2Y_12_ on OXT neurons can also be induced by other types of metabolic challenges. Indeed, the over-representation of the double-positive *Oxt*^+^*P2ry12*^+^ neurons occurs also upon diverse conditions altering energy homeostasis, such as aging or disruption of circadian rhythmicity ^40–42,44–46^, but not upon fasting (**Figure 2C**). Accordingly, we also detected a strong OXT neuron-specific expression of human P2Y_12_ receptor in post-mortem PVH tissues from patients with diabetes mellitus (**Figures 2D, S2C** and **Table S1**). Altogether, we identified that Western diet or hyperglycemia induces neuronal expression of P2Y_12_ receptors which may represent the key regulator of the purinergic pathway in the metabolically stressed hypothalamus.

### Down-regulation of P2Y_12_ signaling protects mice from obesity

Melanocortinergic projections from the first order neurons release α-melanocyte stimulating hormone (α-MSH) in the PVH and other brain regions to induce satiety ^47–49^ via activation of Gα_s_-coupled melanocortin 4 receptor (MC4R) on the second order neurons ^50^. Activation of MC4R or other Gα_s_-associated GPCR receptors on those second order neurons initiates the adenylyl cyclase/cAMP/kinases cascade leading to the expression of Fos proto-oncogene, activator protein 1 (AP-1) transcription factor subunit (or c-Fos), a marker and inducer of neuronal activity ^51–54^. Hence, we hypothesized that Gα_i_-dependent inactivation of the adenylyl cyclase/cAMP/c-Fos axis downstream of P2Y_12_ antagonizes the Gα_s_ signaling in the second order neurons, for example, via MC4R, which might lead to obesity (**Figure 3A**).

**Figure 3.**
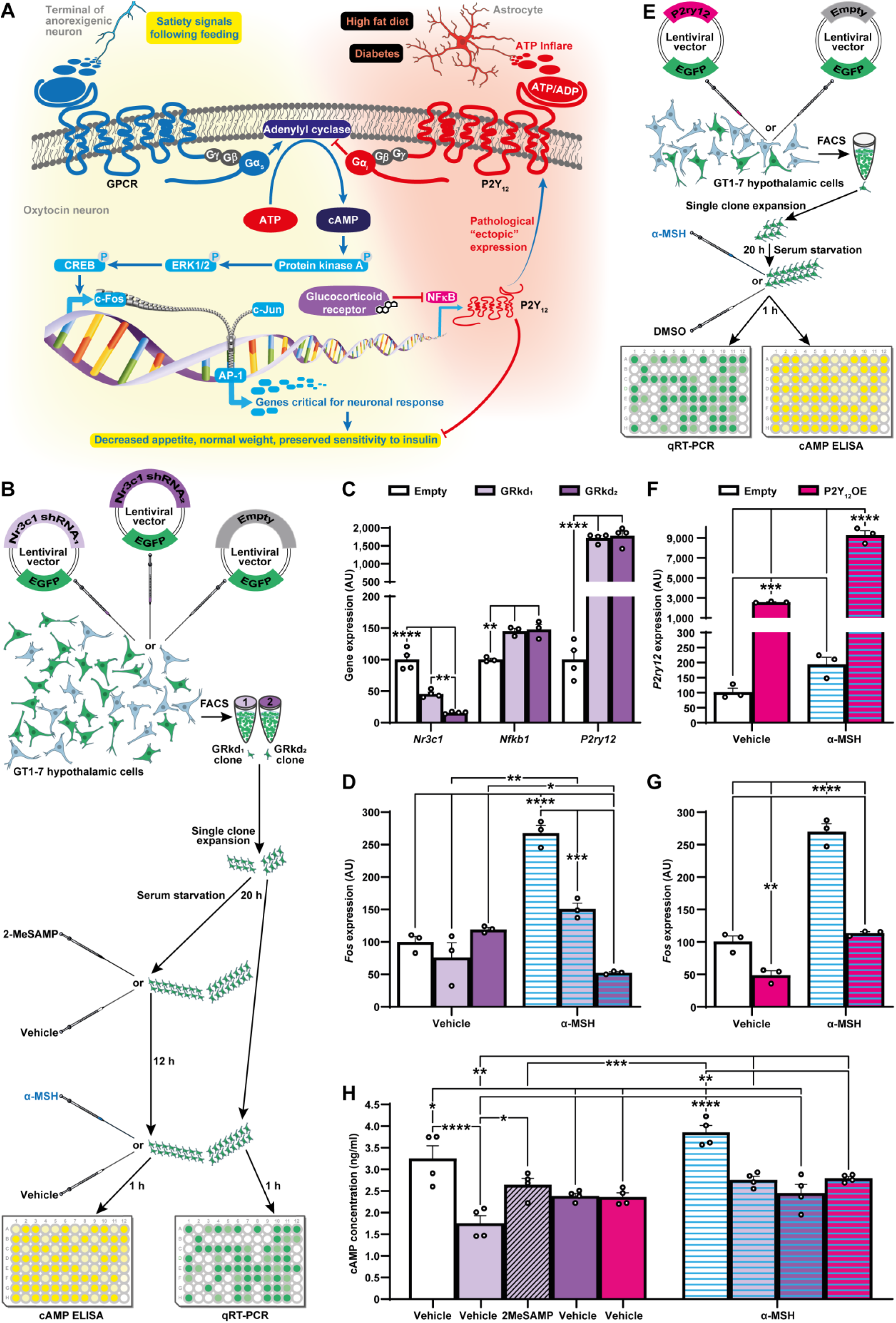
P2Y_12_ diminishes responsiveness of hypothalamic cells. (**A**) Schematic representation of the hypothetic mechanism involving purinergic pathway interactions within PVH^OXT^ neurons. (**B-D**) Scheme of the experiment to transduce GT1-7 hypothalamic cells with lentiviral vectors expressing *Nr3c1* small hairpin RNA (shRNA) (**B**) and qRT-PCR analysis of *Nr3c1*, *Nfkb1*, *P2ry12* (**C**, n=4) and *Fos* (**D**, n=3) genes with or without 1 h-stimulation by 100 nM α-melanocyte-stimulating hormone (α-MSH). (**E-G**) Experimental design to transduce GT1-7 hypothalamic cells with a lentiviral vector overexpressing *P2ry12* followed by α-MSH-mediated stimulation (**E**) and qRT-PCR analysis of *P2ry12* (**F**) or *Fos* (**G**) expression (n = 3). (**H**) ELISA-based analysis of cyclic adenosine monophosphate (cAMP) levels in GT1-7 hypothalamic cells transduced by lentiviral vectors expressing *Nr3c1* shRNA or *P2ry12*, with or without 12 h-pretreatment by P2Y_12_ inhibitor 2-methylthio-AMP (2MeSAMP) followed by 1 h-incubation with 100 nM α-MSH (n = 4). Error bars represent SEM. *, *p* < 0.05; **, *p* < 0.01; ***, *p* < 0.001; ****, *p* < 0.0001 as analyzed by one-way ANOVA followed by post-hoc Holm-Šídák’s test. See also **Figure S3**.

To induce P2Y_12_ expression *in vitro*, we first aimed to alter the expression of the upstream glucocorticoid receptor (GR)/NF-κB axis. Recently, we discovered that knockout of GR in developing OXT or mature PVH^OXT^ neurons leads to P2Y_12_ overexpression-dependent obesity, which can be aggravated by HFD treatment (LY, AB, SSN, DL, SH, YZ, IAV in preparation). Similar to our *in vivo* knock-out data, knock-down of GR in hypothalamic GT1-7 cells expressing MC4R ^55,56^ led to a drastic upregulation in *P2ry12* expression, aided by an up-regulation of its direct activator NF-κB (**Figure 3B,C**). Furthermore, down-regulation of GR prevents activation of *Fos* by α-MSH (**Figure 3D**). The latter neuropeptide, notably, also induces GR expression, which might contribute to preserved responsiveness of these cells as a positive feedback loop (**Figure S3A**). To directly prove *in vitro* the role of P2Y_12_ in diminished responsiveness to α-MSH, we next overexpressed this purinergic receptor, and, accordingly, detected a diminished expression of *Fos* and inability of α-MSH to induce its transcription (**Figure 3E-G**). These results are in agreement with the reduced levels of cAMP, an upstream regulator of c-Fos, in response to P2Y_12_ overexpression or GR knockdown (**Figure 3H**). Interestingly, the effect of GR knockdown on cAMP was attenuated by application of 2-methylthio-AMP (2MeSAMP), a P2Y_12_ inhibitor, further strengthening the contribution of this receptor in the loss of responsiveness upon GR down-regulation.

Recently we demonstrated that knock-out of P2Y_12_ in PVH^OXT^ neurons attenuates weight gain and fat accumulation in mice with GR deletion in these neurons (LY, AB, SSN, DL, SH, YZ, IAV in preparation). We next sought to further validate the detrimental role of P2Y_12_ in PVH^OXT^ neurons in the HFD model, which represents a more disease-relevant pathological condition. Similarly, PVH^OXT^ neuron-restricted knockout of P2Y_12_ successfully prevented diet-induced weight gain (**Figures 4A,B** and **S3B**), further confirming the causative pathophysiological role of PVH^OXT^ neuron-expressed P2Y_12_ receptors in metabolic phenotypes.

**Figure 4.**
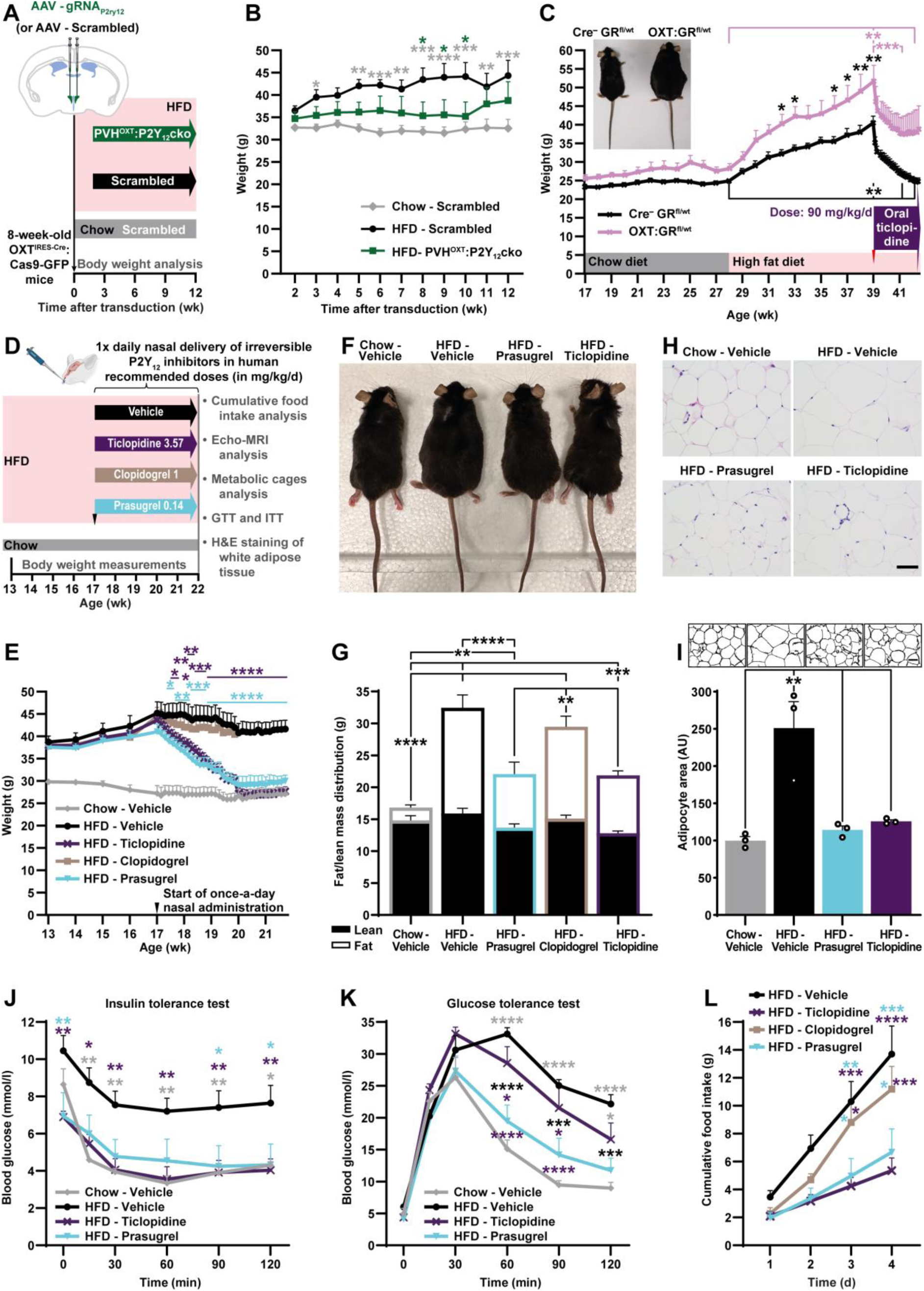
Down-regulation of P2Y_12_ signaling attenuates obesity in mice. (**A**,**B**) Experimental design (**A**) and weight (**B**) in male HFD-fed mice with P2Y_12_ knock-out in PVH^OXT^ neurons and Scrambled guide RNA (gRNA) recombinant adeno-associated viral vector (rAAV)-injected controls fed with chow or HFD (n = 6, 8, 7, respectively). (**C**) Representative macrophotograph on Week 39 (**top left**) and weight of HFD-fed female OXT:GR^fl/wt^ or Cre-deficient GR^fl/wt^ littermates receiving 90 mg/kg/d of prodrug of blood brain barrier-crossing irreversible P2Y_12_ inhibitor ticlopidine via drinking water (n = 4). (**A-I**) Experimental design (**D**), weight (**E**), representative macrophotograph (**F**), Echo-MRI-based fat/lean mass distribution analysis on Week 20 (**G**), representative microphotographs (**H**) with adipocyte edges detection and area quantification of hematoxylin and eosin-stained perigonadal white adipose tissue (**I**, n = 3), insulin (**J**) and glucose (**K**) tolerance tests, cumulative food intake on Week 20 (**L**) in HFD-fed male mice nasally administered once a day with 1 mg/kg clopidogrel, 0.14 mg/kg prasugrel, 3.57 mg/kg ticlopidine or the same volume of the vehicle and chow diet-fed male mice receiving vehicle (unless otherwise stated, n = 5, 5, 7, 5, and 5, respectively, except n = 6 for the HFD-Ticlopidine group in tolerance tests depicted in panels **J** and **K**). Error bars represent SEM. *, *p* < 0.05; **, *p* < 0.01; ***, *p* < 0.001; ****, *p* < 0.0001 as analyzed by two-way ANOVA with post-hoc Holm-Šídák’s test of HFD-Ticlopidine and HFD-Prasugrel vs. HFD-Vehicle groups (**E**) or between all groups (B,**J-L**), one-way ANOVA followed by post-hoc Holm-Šídák’s test (**G**,**I**) or two-way ANOVA with Holm-Šídák’s test followed by repeated measurements ANOVA within each group with Holm-Šídák’s test of 3 time points vs. the peak of the phenotype prior to the treatment start at Week 39 (**C**). Significant differences are depicted over the datapoint which is compared vs. respective groups outlined with color; for fat/lean distribution, they depict differences in fat content. Scale bars: 50 µm. See also **Figures S3** and **S4**.

With the objective to further explore *in vivo* the contribution of elevated P2Y_12_ signaling in metabolic regulation and obesity phenotype by a complementary, pharmacological approach, we orally administered HFD-fed OXT:GR^fl/wt^ or GR^fl/wt^ littermates (**Figure 4C**) with 90 mg/kg/day dose ^57^ of ticlopidine, a prodrug of an irreversible P2Y_12_ inhibitor, which is capable to cross blood-brain barrier. Indeed, such an excessive dose (27 times the human daily recommended dose) of ticlopidine rapidly attenuated the diet-induced obesity in OXT:GR^fl/wt^ mice and normalized weight in the control group, in agreement with the loss-of-function experiment shown above, demonstrating that purinergic signaling contributes to the progression of this metabolic phenotype (**Figure 3C** and **Table 1**). Notably, we failed to rescue the obesity phenotype in HFD mice by high doses (27 times the human daily recommended dose) of other irreversible P2Y_12_ inhibitor prodrugs, such as clopidogrel and prasugrel, or by lower concentrations (9 times the human daily recommended dose) of ticlopidine (**Figure S3C**).

**Table 1.** Summary of metabolic activities analyzed for P2Y_12_ inhibitors.

| P2Y<sub>12</sub> inhibitor | Route, sex | Dose (mg/kg/d) | Reduction |  |  |  |  |
| --- | --- | --- | --- | --- | --- | --- | --- |
|  |  |  | Weight (maximal)<sup>†</sup> | Food intake<sup>‡</sup> | GTT AUC<sup>‡</sup> | ITT AUC<sup>‡</sup> | Fasting glucose<sup>‡</sup> |
| Ticlopidine | Oral, female | 90 | 27.41 ± 5.30% after 3.4 wk, *p* = 0.00575 | Not analyzed | Not analyzed | Not analyzed | Not analyzed |
|  | Nasal, male | 3.57 | 38.27 ± 2.39% after 3.1 wk, *p* = 0.00019 | 56.89 ± 14.4%, *p* = 0.00294 | 8.31 ± 11.0%, *p* = 0.4615 | 86.93 ± 22.5%, *p* = 0.0039 | 34.03 ± 8.18%, *p* = 0.0181; 6.9 ± 0.3 |
| Prasugrel | Nasal, male | 0.14 | 28.83 ± 6.1% after 3.1 wk, *p* = 0.0103 | 51.05 ± 15.6%, *p* = 0.00553 | 37.56 ± 11.4%, *p* = 0.0118 | 78.35 ± 23.5%, *p* = 0.00537 | 33.65 ± 14.36%, *p* = 0.0181; 6.94 ± 1.26 |
| Ticagrelor | Nasal, male | 2.40 | 20.73 ± 4.84% after 2.7 wk, *p* = 0.00793 | 64.29 ± 13.1%, *p* = 0.00023 | 10.4 ± 9.4%, *p* = 0.2923 | 54.47 ± 19.6%, *p* = 0.0172 | 29.2 ± 9.95%, *p* = 0.0252; 6.54 ± 0.28 |
| Cangrelor | Nasal, male | 0.18 | 24.49 ± 2.21% after 2.7 wk, *p* = 0.00018 | 75.30 ± 13.1%, *p* < 0.0001 | 34.7 ± 9.4%, *p* = 0.00636 | 61.43 ± 19.6%, *p* = 0.0172 | 31.6 ± 12.87%, *p* = 0.0252; 6.32 ± 0.8 |

<sup>†</sup> Maximal body weight loss detected in this study, with the differences before vs. after drug administration in wild-type mice with diet-induced obesity ± standard error of means (SEM), statistically analyzed by paired two-tailed Student's *t*-test (from experiments depicted in **Figures 4C,EB** and, for cangrelor and ticagrelor, **5B**). Notably, the latter two drugs were first administered with half a human daily dose for periods indicated in **Figure 5A**.
<sup>‡</sup> Reduction of the area under the curve (AUC) of cumulative food intake, glucose (GTT) or insulin tolerance tests (ITT) and 4 h-fasting blood glucose levels (in mmol/l) vs. the HFD-Vehicle groups in the experiments depicted in **Figures 4J-L** and **5E-G** ± SEM, statistically analyzed by one-way ANOVA followed by Holm-Šidák's post hoc test. Baseline of 0 g and 4 mmol/l was used for food intake and tolerance tests, respectively. Additionally, absolute 4 h-fasting glucose levels are depicted.

### Nasal administration of P2Y_12_ inhibitors in clinically relevant dosages normalizes weight and food intake in mice

To lower the administered doses of P2Y_12_ inhibitors to the levels that are recommended in humans for daily peroral administration, while maintaining high concentrations in the brain, we conceptualized a nasal way of delivery (**Figure 4D**). Rewardingly, once-a-day nasal administration of irreversible inhibitor prodrugs ticlopidine, prasugrel, but not vehicle or clopidogrel, in clinically relevant peroral human doses validated for safety (**Figure S3D**), rapidly reversed obesity and attenuated fat mass accumulation in HFD-fed mice (**Figure 4E-G** and **Table 1**). Moreover, after normalization, the weights did not continue falling upon treatment of the inhibitors (**Figure 4E**), similar to the absence of weight loss in chow-fed animals (**Figure S3D**), suggesting that the anti-obesity effect is dependent on the metabolic state and is not the result of toxicity. Notably, the anti-obesity effect of ticlopidine was lost upon further halving of the dose (**Figure S3E**), indicating that the human daily recommended peroral dose approaches to the minimal effective concentration in mice. Both ticlopidine and prasugrel normalized HFD-induced adipocyte hypertrophy, fasting glucose levels and the performance in the insulin tolerance test (**Figure 4H-J** and **Table 1**). Interestingly, prasugrel, but not ticlopidine, normalized the performance in the glucose tolerance test (**Figure 4K** and **Table 1**). Mechanistically the anti-obesity effects of the P2Y_12_ inhibitors are mostly attributed to the reduction in food intake (**Figure 4L**), with no detected effect in locomotor activity or restoration of energy expenditure (**Figure S4**).

With the goal to identify the lowest effective dose, we halved the human daily recommended doses also for prasugrel, as well as for the two reversible P2Y_12_ inhibitors cangrelor and ticagrelor (**Figure 5A**). Interestingly, similar to irreversible inhibitor prodrugs, both cangrelor and ticagrelor demonstrated a strong anti-obesity potential at clinically relevant doses when administered nasally (**Figures 5B-C, S5A** and **Table 1**), with cangrelor being the only P2Y_12_ inhibitor demonstrating effectiveness in mice at subhuman concentrations. These results, again, suggest that the human daily recommended peroral doses of prasugrel and ticagrelor approach to the minimal effective concentrations in mice. Importantly, we used nucleic magnetic resonance spectroscopy to verify the abundance of nasally administered ticagrelor and its metabolites in the hypothalamus (**Figure S5B**). Interestingly, even up to ten days after withdrawal of the nasal administration of ticagrelor or cangrelor, the animals revealed no signs of weight regaining (**Figure 5D**). While both cangrelor and ticagrelor normalized food intake, fasting glucose levels and performance at insulin tolerance test, only cangrelor normalized glucose tolerance (**Figure 5E-G** and **Table 1**). Indeed, the latter was significantly attenuated, but not normalized by ticagrelor.

**Figure 5.**
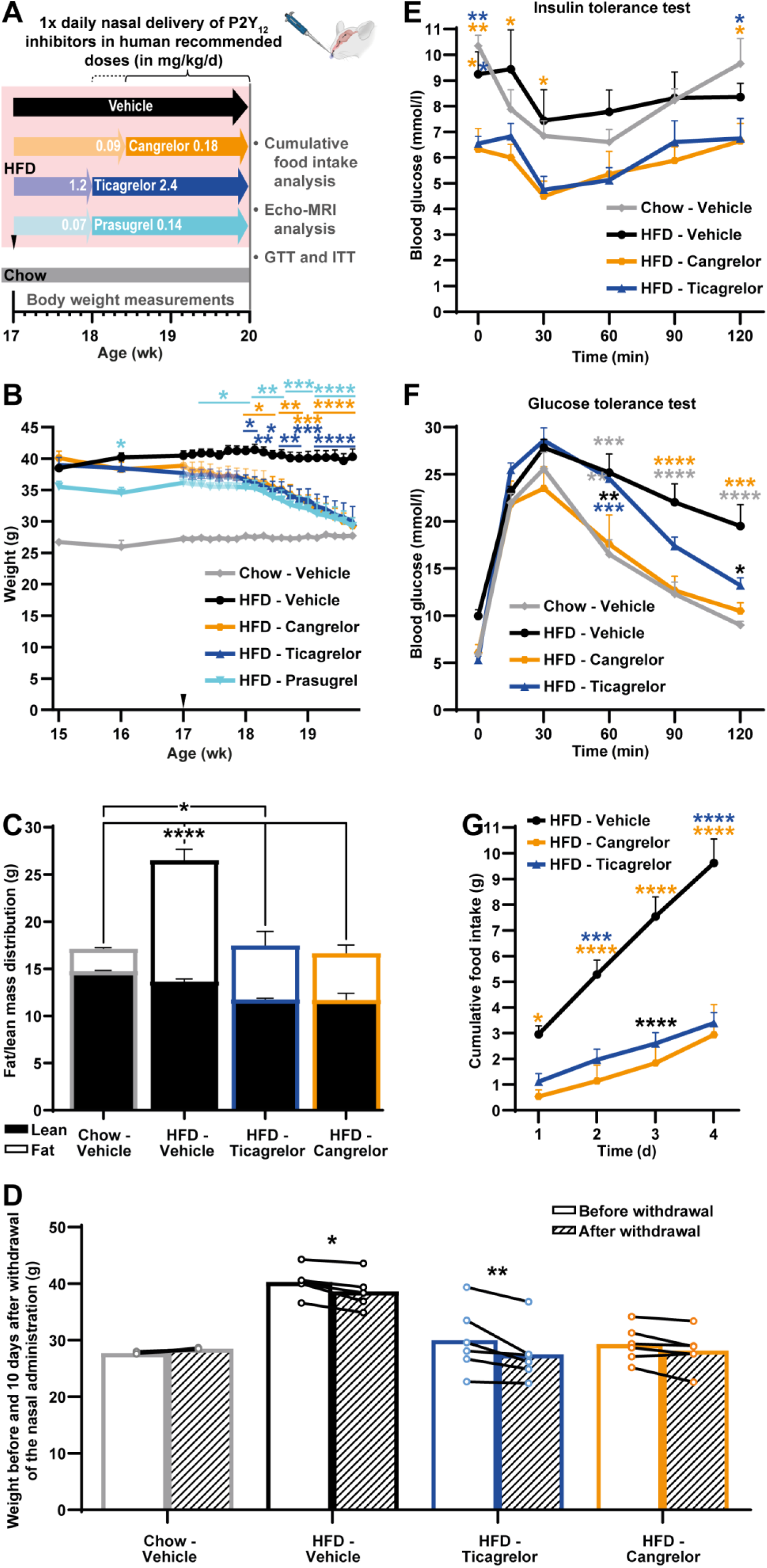
Nasally administered P2Y_12_ inhibitors reverse obesity and hyperphagia in mice. (**A-C**) Experimental design to determine the minimal effective dose of the indicated P2Y_12_ inhibitors (**A**), weight (**B**), Echo-MRI-based fat/lean mass distribution analysis (**C**, n = 5), in chow diet- and HFD-fed male mice nasally administered once a day with vehicle or 0.09 mg/kg cangrelor for 10 days, 1.2 mg/kg ticagrelor and 0.07 mg/kg prasugrel for 1 wk, followed by doubling the doses (n = 5, 5, 6, 6, and 5, respectively, unless otherwise stated). (**D**) Individual weight data in chow diet- and HFD-fed male mice from the above-stated experiment on the last day of nasal administration of vehicle, ticagrelor or cangrelor and ten days after withdrawal (n = 5, 5, 6 and 6, respectively). (**E-G**) Insulin (**E**) and glucose (**F**) tolerance tests (n = 5), cumulative food intake (**G**) in chow diet- and HFD-fed male mice from the above-stated experiment nasally administered once-a-day with vehicle or cangrelor, ticagrelor and prasugrel (n = 5, 5, 5, 6, and 5, respectively, unless otherwise stated). Error bars represent SEM. *, *p* < 0.05; **, *p* < 0.01; ***, *p* < 0.001; ****, *p* < 0.0001 as analyzed by two-way ANOVA with post-hoc Holm-Šídák’s test of HFD-Inhibitors vs. HFD-Vehicle groups (**B**) or between all groups (**E-G**), one-way ANOVA with post-hoc Holm-Šídák’s (**C**) and repeated measurements two-way ANOVA with post-hoc Holm-Šídák’s test (**D**). Significant differences are depicted over the datapoint which is compared vs. respective groups outlined with color; for fat/lean distribution, they depict differences in fat content. See also **Figure S5**.

### Nasal administration of cangrelor spray counteracts obesity in non-human primates

Encouraged by the effect of P2Y_12_ blockers in rodents, we sought to apply them to counteract obesity in crab-eating macaques (*Macaca fascicularis*). In agreement with previous reports ^58^, about 15% of aged in our facility tend to voraciously consume all *ad libitum* provided food, developing obesity. Hence, such animals are subjected to calorie-restricted conditions to prevent age-associated pathologies ^59–61^ (**Table S2**). To resume the weight gain in our experimental design, we doubled the dose of the provided afternoon chow diet portion in overweight macaques while simultaneously administering once-a-day nasal spray with human daily recommended doses of cangrelor or vehicle (**Figure 6A**). In agreement with our results in mice, cangrelor protected macaques from weight gain not only during the nasal administration, but also up to eight weeks after withdrawal of the drug (**Figure 6B-D**), further strengthening the perspective to use the P2Y_12_ inhibitors in the treatment of metabolic disorders.

**Figure 6.**
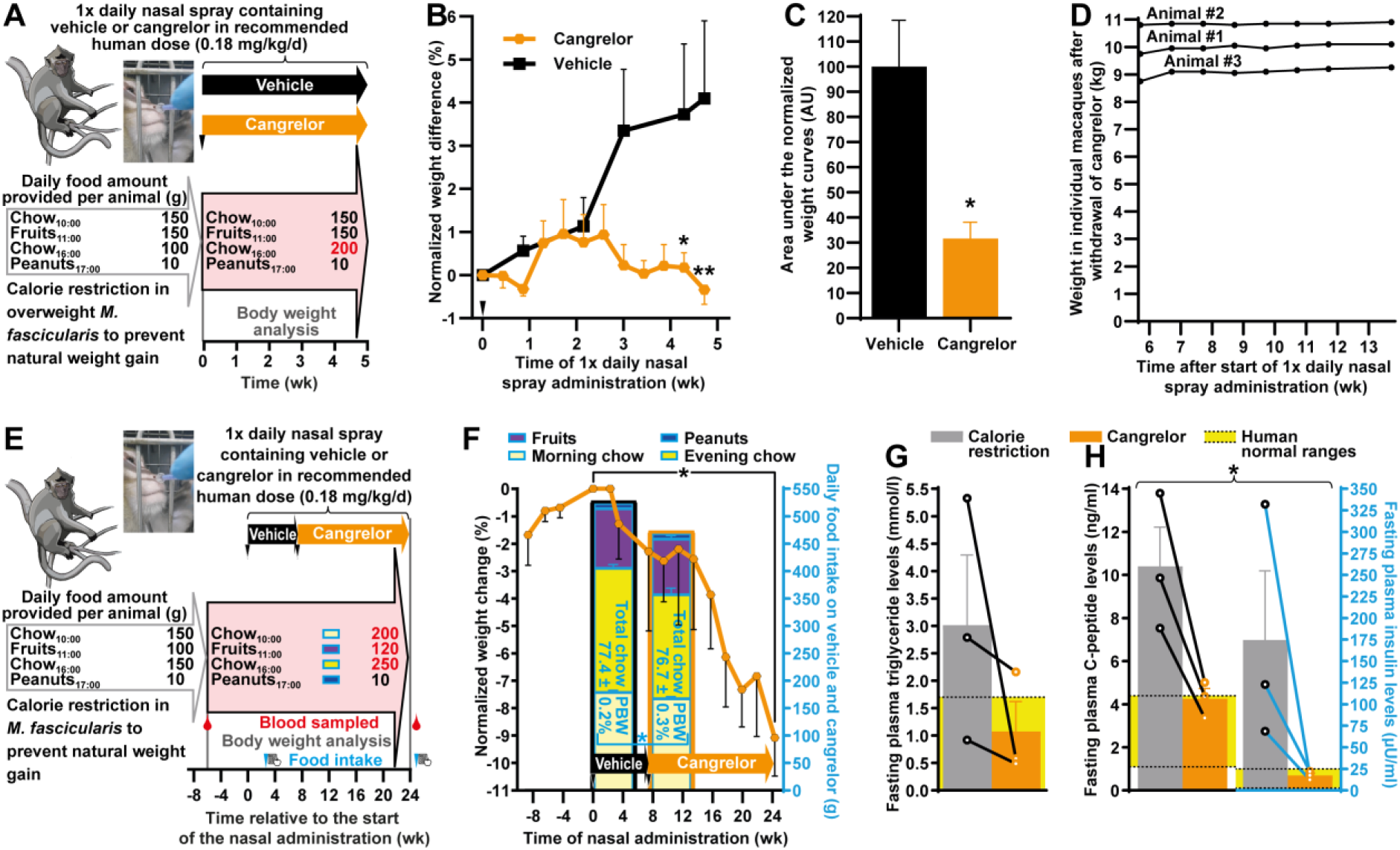
Nasally administered cangrelor counteracts obesity in non-human primates. (**a**-**d**) Experimental design (**A**), normalized weight change (**B**), area under the curve of the normalized weight (**C**) and individual weight data following the drug withdrawal (**D**) in overweight male crab-eating macaques (*Macaca fascicularis*) receiving a doubled portion of afternoon chow diet compared to normal maintenance conditions in parallel with once-a-day nasal administration of spray containing vehicle or 0.18 mg/kg/d cangrelor (n = 3). (**E-H**) Scheme of the long-term nasal administration of 0.18 mg/kg/d cangrelor spray (**E**), normalized weight change (black) overlaid with daily food intake (cyan) composition indicating percentage by weight (PBW) of total chow food consumed per day on the 28^th^ and 145^th^ days after start of vehicle and cangrelor spray administration, respectively (**F**), and fasting plasma levels of triglycerides (**G**), C-peptide and insulin (**H**) on the −97^th^ (last day before the termination of calorie restriction) and 157^th^ (last day before the termination of cangrelor administration) days relatively to the start of cangrelor spray administration in male crab-eating macaques receiving daily amount of chow diet increased by 50% and fruits—by 20% compared to normal maintenance conditions (n = 3). Error bars represent SEM. *, *p* < 0.05; **, *p* < 0.01 as analyzed by two-way ANOVA with post-hoc Holm-Šídák’s test (**B**), one-way repeated measurements ANOVA with post-hoc Holm-Šídák’s test at the end of each nasal treatment (Weeks 7 and 24) vs. Week 0 (weight change in **F**), paired (PBW of chow food in **F**) or unpaired (**C**) two-tailed Student’s t-test, paired Hotelling’s T^2^ test for cangrelor samples of C-peptide and insulin vs. calorie restricted samples from the same animals (**H**). Averaged Cohen’s d for triglycerides, C-peptide and insulin were 1.139, 2.654 and 1.598, respectively. For the area under the curve analysis, the smallest normalized weight value was used as a baseline. Significant differences vs. respective groups are outlined with color.

To lower the body weight in *M. fascicularis* by prolonged daily cangrelor spray administration, we first lifted the calorie restriction in macaques, increasing both the provided amount of fruits and vegetables by 20% and chow diet by 50% (**Figure 6E**). Consistent with the results in mice, cangrelor, but not vehicle administration, significantly reduced weight and percentage by weight of daily consumed chow food in these animals (**Figure 6F**). Previous reports indicated insulin resistance and altered lipid homeostasis in individuals with enhanced body mass index ^62,63^. In agreement, when an analogous index, the body condition score (BCS), was moderately increased in macaques (**Table S2**), they, despite calorie restriction, revealed elevated levels of triglycerides, insulin and C-peptide (**Figure 6G,H**), but not other parameters (**Table S3**) in the fasting plasma. Interestingly, nasal administration of cangrelor maintained triglycerides in a normal range in two out of the three macaques, although the overall down-regulation tendency here did not reach a statistical significance, despite a large effect size (**Figure 6G**). Even larger, but also significant effects were observed for cangrelor spray-dependent down-regulation of C-peptide and insulin levels, where the former again in two out of the three animals and the latter in all animals were restored to the normal ranges (**Figure 6H**). These observations confirm the data in mice that metabolism-alleviating properties of P2Y_12_ inhibitors extend beyond merely weight reduction. Altogether, these results further support an evolutionary conserved role of purinergic signaling in metabolic control and provide a translational perspective for nasal administration of P2Y_12_ inhibitors to treat metabolic diseases.

### Over-expression of P2Y_12_ in PVH^OXT^ neurons is sufficient to drive obesity in mice

To explore the sufficiency of the P2Y_12_ expression to induce obesity, we next overexpressed this receptor specifically in PVH^OXT^ neurons of adult mice by transducing OXT^IRES-Cre^ mice with rAAV vector expressing P2Y_12_ (**Figures 7A** and **S6**). In agreement with our *in vitro* data, the P2Y_12_-overexpressing OXT neurons in the PVH became less responsive to hyperosmotic stress (**Figure 7B,C**), which is known to stimulate OXT neurons ^64^. Moreover, PVH^OXT^:P2Y_12_ mice fed with normal chow diet gained weight, which could be reversed by nasally administered prasugrel in human daily recommended dose (**Figure 7D-E**), with no regain of weight even up to ten days after withdrawal of the drug (**Figure 7F**). Notably, PVH^OXT^:P2Y_12_ mice fed with chow diet also developed insulin resistance and hyperphagia, which could also be reversed by nasal prasugrel administration (**Figure 7G-I**). In agreement with our previous data, a tendency towards a decreased energy expenditure or locomotor activity in P2Y_12_-overeexpressing mice did not reach a statistical significance (**Figure S7**), confirming that the weight gain-promoting effect of the over-activated purinergic signaling is mainly driven by hyperphagia. Accordingly, reversal of obesity by P2Y_12_ inhibitors predominantly relies on their anorexigenic action (**Figures 4L**, **5G** and **7I**), in agreement with studies demonstrating appetite-suppressing effects of OXT ^16,17^. Altogether, hyperphagic obesity and glucose disposal impairment induced by P2Y_12_ expression on PVH^OXT^ neurons can be reversed by the inhibitors of this ADP/ATP receptor, confirming its causative obesogenic role.

**Figure 7.**
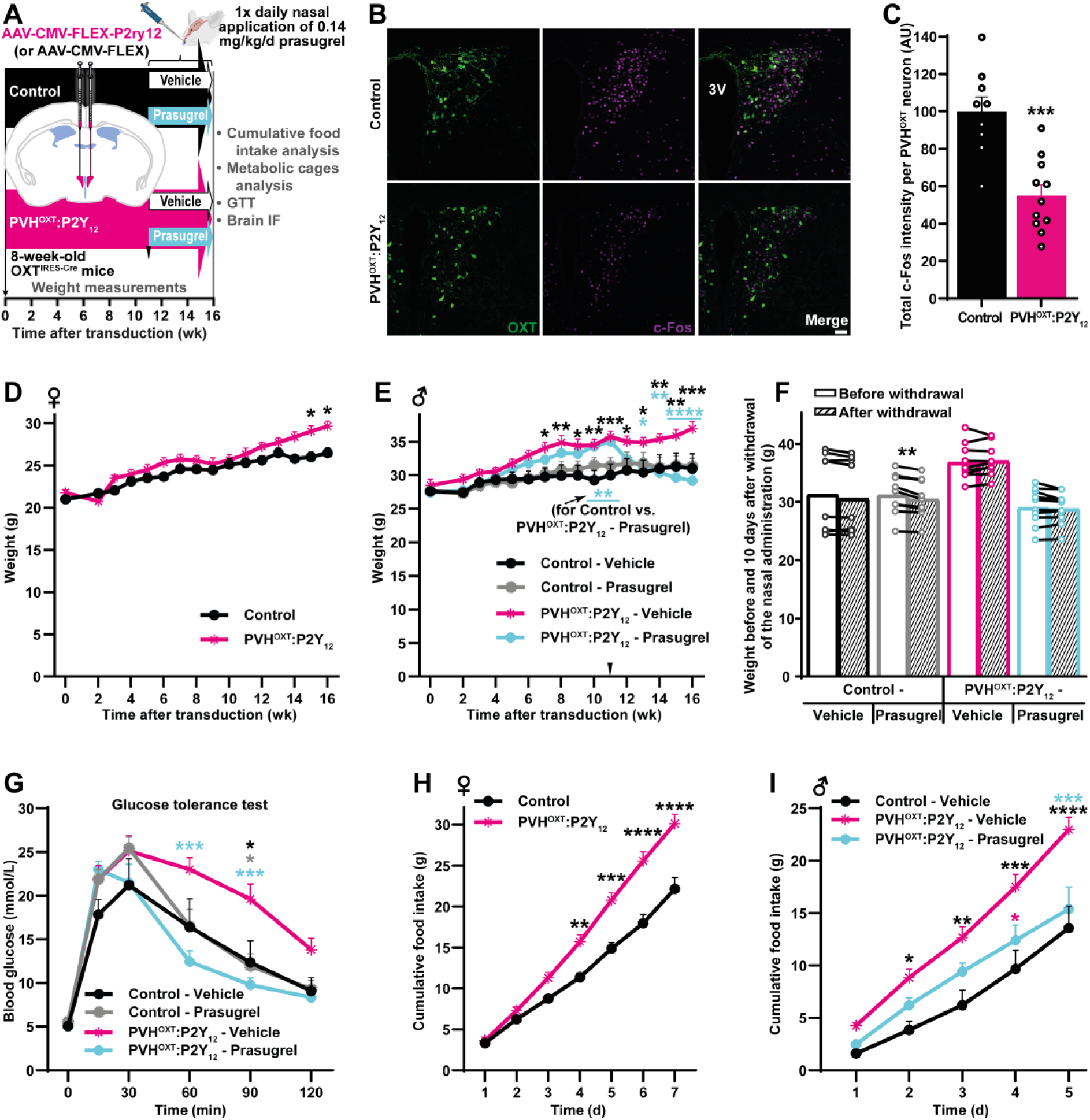
Over-expression of P2Y_12_ in PVH^OXT^ neurons drives hyperphagic obesity and insulin resistance in mice. (**A-C**) Experimental design implementing flip-excision (FLEX) over-expression strategy (**A**), representative microphotographs (**B**) and quantification (**C**) of hyperosmotic stress-induced c-Fos expression in PVH^OXT^ neurons (n = 8 males and 1 female for Controls and 7 males and 4 females for PVH^OXT^:P2Y_12_ mice). (**D-I**) Weight (**D**,**E**), individual weight data before and ten days after withdrawal of the drug (**F**), glucose tolerance test (**G**) and cumulative food intake analyses (**H,I**) in male (**E**,**F,G,I**) and female (**D**,**H**) OXT^IRES-Cre^ mice after transduction of their PVH with the indicated recombinant adeno-associated viral vectors (rAAVs) followed by once-per-day nasal administration of male mice with 0.14 mg/kg/d prasugrel or vehicle. For females n = 7 and 16 for Control and PVH^OXT^:P2Y_12_ groups, respectively; for males n = 10 for the PVH^OXT^:P2Y_12_-Prasugrel group and 9 for all others, except for food intake analysis (**I**) with n = 4. Error bars represent SEM. *, *p* < 0.05; **, *p* < 0.01; ***, *p* < 0.001; ****, *p* < 0.0001 as analyzed by two-way ANOVA with Holm-Šídák’s test between Control-Vehicle, PVH^OXT^:P2Y_12_-Vehicle and PVH^OXT^:P2Y_12_-Prasugrel groups (**E**) or between all groups (**D**,**G-I**), unpaired two-tailed Student’s t-test (**C**) and repeated measurements two-way ANOVA with post-hoc Holm-Šídák’s test (**F**). Significant differences are depicted over the datapoint (unless explicitly indicated) which is compared vs. respective groups outlined with color. Scale bar: 50 µm. See also **Figures S6** and **S7**.

## DISCUSSION

Understanding the metabolic stress-induced signaling changes in the hypothalamic circuits is key to develop effective therapeutic strategies. Our results indicate HFD- or hyperglycemia-evoked stimulation and profound reorganization of purinergic signaling in the PVH by linking astrocytes and OXT neurons within the pathological signaling loop. The gradual time-dependent frequency elevation of ATP release from astrocytes in the PVH of HFD-exposed mice suggests that the extent of metabolic damage is encoded by populational Inflares, in agreement with our previous data ^23^. Decoding of these ATP signals by overexpressed P2Y_12_ receptors on PVH^OXT^ neurons leads to hyperphagic obesity and insulin resistance, which can be attenuated by nasal administration of human recommended daily doses of inhibitors of this ATP/ADP receptor.

HFD-induced Inflares share identical releasing mechanism with other injury modalities ^23^, which was validated by pharmacological inhibition and astrocyte-specific knockout of pannexin 1. Interestingly, while Inflares exhibit similar single-event properties across different injury intensities, diabetic mice revealed prolonged duration accompanied with the elevated frequency, suggesting that different modalities of purinergic signals may encode specific metabolic challenges, which could be due to differences either in the upstream signaling of pannexin 1 within astrocytes or in extracellular mechanisms modulating ATP levels. As our experiments are conducted in the acute brain slice system, we may not exclude the role of ATP events caused by local cell apoptosis during the sample preparation or *ex vivo* imaging process. Future approaches using gradient refractive index lens-based *in vivo* imaging in the hypothalamus will be able to characterize the continuous changes of purinergic signals in the native environments.

Previously we reported that due to the abundance of purinergic receptors, the information encoded by Inflares can be efficiently retrieved and further transduced by microglia as the key decoder ^23^, which may modulate their migration, proliferation and initiation of inflammatory responses leading to reactive microgliosis ^65,66^, also detected by us in the PVH from patients with diabetes mellitus. Here we identified that in addition to microglia, OXT neurons may be context-dependently converted to yet another decoder, resulting in aggravation of pathological phenotypes. Importantly, the HFD-induced P2Y_12_ expression on OXT neurons emerges prior to the onset of weight gain. This evolutionary conserved context-dependent expression may be related to the duration or extent of metabolic phenotypes. Indeed, among PVH^OXT^ neurons from patients with diabetes mellitus, we detected various numbers of cells strongly positive for P2Y_12_. OXT (and possibly other) neurons might react to metabolic stressors or other injuries with different timing and/or dynamics, leading to alterations of the purinergic signaling and reshaping the neuronal circuits in healthy and diseased brain. Overall, apart from the pathological role in exacerbation of the metabolic phenotypes, we presume that this purinergic circuit may contribute to other physiological functions, while the epigenetic landscape of these neurons might be permissive for other stress-related contexts or not yet explored conditions, for example, related to circadian clock or aging, to promote the emergence of P2Y_12_-dependent (patho)physiological modalities of OXT (and possibly other types of) neurons. Moreover, potentially, P2Y_12_-expressing neurons may also sense ATP from other sources, for example, systemic circulation ^67^. However, the data from Woo-ping Ge and colleagues demonstrate 3.9- and 6.63-fold decreased hypothalamic ATP abundance in young P56 and old P900 mice, respectively, compared to the postnatal day P4, as well as a 1.29-fold decreased PVH abundance in P900 vs. P56 mice. In contrast, the abundance of the ATP precursor adenosine, which is fully penetrable through the blood-brain barrier, declines by only 1.57- and, not significantly, by 0.96-fold within P56 and P900 hypothalami, respectively, compared to P4, while not significantly increasing by 1.17-fold within the PVH upon aging, suggesting the relevance of the local ATP production and release within this region (WG et al., in preparation).

In this work, we used the loss-of-function model of GR, the negative regulator of P2Y_12_, with knock-out onset during embryonic development. We and others identified abundant expression of GRs on PVH^OXT^ neurons ^68^, which are responsible for both suppression of appetite and stimulation of energy expenditure ^14–17^ and receive projections from various neurons regulating energy homeostasis, including the first order melanocortinergic neurons ^8–11,13^. In contrast to the HFD-induced PVH^OXT^ neuron-restricted double-knockout strategy that we recently implemented (LY, AB, SSN, DL, SH, YZ, IAV in preparation) the isolated— without a use of HFD or additional gene manipulations—loss or gain of function of P2Y_12_, which result in reciprocal metabolic phenotypes, are functionally more valuable to establish the causative pathophysiological role of this receptor in these neurons.

In agreement with our previous work on knock-out models of GR ^36^, isolated loss of GR in GT1-7 cells does not lead to *Fos* down-regulation per se, suggesting that inhibition of the NF-κB/P2Y_12_ axis by GR becomes especially critical only upon anorexigenic stimulation of neurons, for example, by α-MSH. Our *in vitro* data on attenuation of the responsiveness of hypothalamic cells by P2Y_12_ overexpression was validated by the gain-of-function Cre-LoxP-FLEX model. Indeed, hyperosmotic stress-stimulated c-Fos expression was markedly down-regulated in PVH^OXT^ neurons of PVH^OXT^:P2Y_12_ mice. By no means, however, may we assume melanocortinergic stimulation as the only or primary candidate modality to induce the above-mentioned anorexigenic effects in PVH^OXT^ neurons, especially since only a relatively small subset of these neurons expresses MC4R ^11,69,70^. It remains to be directly demonstrated whether P2Y_12_-assisted inhibition of adenylyl cyclase can counteract Gα_s_-coupled GPCR receptors, such as β-adrenergic receptors, MC4R, receptors of GLP-1, GIP, amylin, BRINP2-related peptide or glucagon. Here, we demonstrated an obesogenic sufficiency of the P2Y_12_ overexpression on PVH^OXT^ neurons, since both hyperphagic obesity and impaired glucose tolerance could be induced even without high calorie diets. Further support of the causative or at least contributing obesogenic role of P2Y_12_ comes from the loss-of-function models demonstrating that knock-out of P2Y_12_ specifically in PVH^OXT^ neurons can attenuate obesity. Yet another confirmation of the causation or contribution of P2Y_12_ comes from the rapid and highly reproducible effect of multiple inhibitors of this ATP/ADP receptor.

Notably, when delivered in extremely high concentrations exceeding the human daily dose by 40 times, clopidogrel, a prodrug of the irreversible P2Y_12_ inhibitor, was reported to significantly reduce body weight and recovery rates in the stroke mouse model, while a phase IV FDA-approved clinical study demonstrated unintentional weight loss in 3528 patients (2.5%) receiving this inhibitor ^71^. Our experiments demonstrated that perorally administered 27 times the daily recommended human dose of another prodrug ticlopidine, but not clopidogrel, was able to counteract obesity. Centrally acting pharmaceuticals represent a promising alternative, but are required to effectively cross the blood-brain barrier when delivered systemically ^72^. With the advantage of the unique anatomical and physiological connection between the nasal cavity and the brain ^73^, nasal delivery of pharmaceuticals emerges as a potential solution to both bypass the blood-brain barrier and mitigate peripheral side effects ^74,75^. Consistently, the metabolic symptoms were effectively reversed not only by nasal administration of clinically relevant doses of direct reversible inhibitors ticagrelor and cangrelor, but also prodrugs ticlopidine and prasugrel, which were plausibly converted to irreversible P2Y_12_ inhibitors by the brain-derived cytochrome P450. Although expressed at much lower levels than in the liver, this enzyme is abundant in the frontal cortex and the hypothalamus, particularly in astrocytes ^76–78^. The successful biodistribution of nasally administered P2Y_12_ inhibitors within the hypothalamus opens a perspective for this route of delivery of these pharmaceuticals to alleviate the metabolic symptoms. Notably, in the insulin tolerance test, the truly meaningful difference between groups with and without P2Y_12_ inhibitors lies in the baseline glucose levels, which is lower in the drug-treated groups. It is worth mentioning that nasal administration of P2Y_12_ inhibitors reversed obesity, but resulted in no further weight reduction beyond the normal weight either in chow or HFD-fed animals, while also induced no weight gain even after withdrawal of the drugs. These results suggest that the observed anti-obesity effect is not merely a consequence of appetite blockade, but rather remodels and alleviates dysfunctional hypothalamic circuits that contribute to the excessive weight gain. Indeed, P2Y_12_ inhibitors remain to be thoroughly studied not only in terms of their pharmacokinetics and pharmacodynamics, but also for their global effects on the organism. For example, the reduction of locomotor activity by prasugrel during the day time could be due to a decreased drive for food in the resting period. On the other hand, the tendency of this inhibitor to alleviate energy expenditure might be an additional contributing factor at certain time points of nasal administration or to be context-dependent.

Strikingly, cangrelor in a form of nasal spray tends to counteract dyslipidemia and is effective to reverse age-associated weight gain and insulin resistance in non-human primates. Our results indicate that at least in certain conditions these drugs may be more effective in metabolic rebalancing than the well-established non-pharmacological interventions, such as calorie restriction, opening the possibility to explore the application of P2Y_12_ inhibitors in humans. Indeed, since OXT neurons of patients suffering from diabetes mellitus start expressing this ATP/ADP receptor, which per se is capable to drive hyperphagic obesity and insulin resistance, pharmacological inhibition of P2Y_12_ might relieve these metabolic symptoms. Nevertheless, we may not rule out that anti-obesity effects of P2Y_12_ inhibitors rely at least in part on other cell populations, including microglia, which may also respond to metabolic context-induced Inflares and indirectly alter anorexigenic circuits within the hypothalamus, especially in the view that some populations of microglia can express OXT ^79,80^. Notably, both our morphological (cell shape and size) and molecular (Iba-1 co-expression) analyses excluded the microglial identity of the emerging OXT^+^P2Y_12_^+^ double-positive cells at any studied time points in response to HFD or diabetes mellitus. Delineating the selective or synergistic effects of purinergic signaling connecting multiple cell types in the hypothalamus is worth further exploration.

Indeed, the increase of targeting range of anti-obesity therapeutics—from activation of a single incretin receptor GLP1 predominantly in the hindbrain (with the body weight loss of about 17%), simultaneous activation of both GLP1 and glucagon (~18%) or GLP1 and GIP (~22.5%) receptors, to triple-agonists (~24%)—gradually potentiates the efficiency while reducing the side effects of the drugs ^2^. Thus, it remains to be demonstrated whether the combination of the aforementioned agonists with the inhibition of yet another, hypothalamic target, P2Y_12_ (loss of the body weight in mice for various P2Y_12_ inhibitors ranging from 20 to 38%, food intake—from 51 to 75%, fasting glucose—from 29 to 34%), constitutes a valuable therapeutic strategy.

In conclusion, we have demonstrated that ATP Inflares and pathological expression of P2Y_12_ on PVH^OXT^ neurons can be induced by HFD or hyperglycemia, while nasal administration of P2Y_12_ blockers can rapidly reverse diet-induced obesity, outlining a potential for these widely used pharmaceuticals as therapeutics to combat metabolic diseases.

## METHODS

### Human samples

Human brain tissue samples used in this study were obtained following ethical guidelines and regulations and approved by the Institutional Review Board of Fudan University, ensuring compliance with the principles outlined in the Declaration of Helsinki. Informed consent was obtained from all donors or their legal representatives prior to sample collection. All patient data were anonymized to protect confidentiality, and tissue handling was conducted according to institutional and national bioethical standards.

### Animals

All experimental procedures in mice (see figure legends for indication of sex for each experiment) were conducted on the C57BL/6 genetic background (at least nine backcrosses) at Shanghai Jiao Tong University and Chinese Institute for Brain Research in accordance with international standards, as approved by the institutional and local authorities of China. *Panx1*^flox/flox^ mice (GemPharmatech, Strain NO. T008074) were used for astrocyte-specific knockout of pannexin 1 and subsequent visualization of ATP events ^81^. B6;129S-*Oxt^tm1.1(cre)Dolsn^*/J (referred to as OXT^IRES-Cre^) mice ^82^ and the tdTomato reporter line B6.Cg-*Gt(ROSA)26Sor^tm14(CAG-tdTomato)Hze^*/J ^83^ (Jackson laboratory, 024234 and 007914, respectively) were crossed to visualize OXT neurons. OXT^IRES-Cre^ and B6;129-*Gt(ROSA)26Sor^tm1(CAG-cas9*,-EGFP)Fezh^*/J ^84^ (Jackson laboratory, 024857) lines were crossed to produce the OXT^IRES-Cre^:Cas-GFP line (homozygous for Cas9-GFP), which then was used for generation of knock-out mutants. OXT^IRES-Cre^ and B6.Cg-*Nr3c1^tm1.1Jda^*/J ^85^ (Jackson laboratory, 021021) lines were crossed to produce the OXT:GR^fl/wt^ mice. Tissue digestion protocol before fluorescence-activated cell sorting (FACS) was followed as described elsewhere ^86^ (see also **Extended Materials and Methods** for more details).

Experiments on cynomolgus (or crab-eating) macaques (*Macaca fascicularis*) were conducted in accordance with the guidelines of the National Institute of Health’s Guide for the Care and Use of Laboratory Animals and approved by the Ethics Committee of Kunming Medical University. Two cohorts of six and 3 obese male macaques aged between 8 to 16 years with body weights ranging from 7.2 to 10.85 kg were individually housed in cages with dimensions of 0.74 m x 0.71 m x 0.74 m, under standard environmental conditions, at 25 ± 2°C, a 14-hour light/10-hour dark cycle, and a relative humidity of 60% with ad libitum access to water (for further details, please refer to **Table S2** and **Extended Materials and Methods**).

### *In vivo* administration of compounds

For peroral administration with drinking water and/or nasal delivery, ticlopidine, clopidogrel, prasugrel hydrochloride, ticagrelor (all from APExBIO; B2164, A5183, B1283, B2166, respectively) and cangrelor tetrasodium (MedChemExpress, 163706-36-3) were used at concentrations indicated in the figures. For the vehicle groups, normal drinking water was used in experiments with peroral administration, 5.4% DMSO (Sigma-Aldrich, D2650) in 0.9% NaCl for nasal administration of ticagrelor or 0.9% NaCl for all other inhibitors. See the Figure legends and **Extended Materials and Methods** for details about numbers ^87^ and sexes ^88–90^ used in each experiment). For nasal delivery in mice according to ^75^, 1-2 µl at a time were alternately and slowly delivered to the entrance of each nostril (total volume of up to 20 µl, depending on the weight of the animal, was applied within 1-5 min) to aid the uptake from the nasal sinus through the sifter plate into the olfactory bulb and spread to the mouse brain. 700 MHz nuclear magnetic resonance (NMR) AVANCE NEO spectrometer (Bruker) was used to analyze the abundance of ticagrelor within the hypothalamus. A spray with cangrelor or vehicle in a total volume of 50 µl per animal once per day was administered nasally to cynomolgus monkeys. For salt loading experiments, 90 minutes after hyperosmotic stimulation of OXT neurons by intraperitoneal injection of 15 ml/kg 1.5 M NaCl ^91^, the isolated brains were fixed in 4% paraformaldehyde (PFA) for subsequent analyses.

### Statistical analysis

Statistical analysis was performed in GraphPad Prism or Python using Shapiro-Wilk normality test, Kruskal-Wallis test followed by Dunn’s multiple comparisons test, two-tailed paired or unpaired Student’s t-tests, one- or two-way ANOVA followed by Holm-Šídák’s post-hoc tests against all groups, unless otherwise stated (see figure legends for details), paired Hotelling’s T^2^ test. Effect sizes were determined by Cohen’s d or, for paired comparisons, by averaged Cohen’s d. Group numbers indicate biological replicates. Results are presented as means ± standard errors of means (SEM). *p*-values of less than 0.05 were considered statistically significant: *, *p* < 0.05; **, *p* < 0.01; ***, *p*<0.001; ****, *p*<0.0001.

## ACKNOWLEDGMENTS

The authors gratefully acknowledge Woo-ping Ge for the data on hypothalamic uptake of ATP from the systemic circulation, Haixia Jiang for help with cell sorting, Jieli Wu for NMR analysis, Tatyana Grinenko for assistance with image analysis, Evgeniya Krasilnikova for insights into clinical use of P2Y_12_ inhibitors, Hua Yang for help with single cell sequencing analysis. We very much thank Hui Lu, Juliang Chen and Honglei Xiao for sectioning of and Anna Berliand for anatomic insights in human brain tissues; Tiemin Liu for invaluable discussions and assistance with obtaining expression data from patients and non-human primates, Vladimir Popov for the technical details of nasal administration; Yan Zhang, Xizhi Guo, Weidong Li, Cheng Hu and Valery Grinevich for the support; Nicola Murgia and Chaweewan Sirakawin for the help with animal management. DNA visualization in the cartoon is from vecteezy.com. This work was funded by the Special project of the Ministry of Life Sciences and the Ministry of Medical Sciences 32241020 and Research Fund for International Scientists 32350610254 from the National Natural Science Foundation of China (NSFC) to IAV; 32371150 from NSFC, the National Key Research and Development Program 2021YFE0116400 from the Ministry of Science and Technology of the People’s Republic of China (MOST), the STI2030-Major project 2021ZD0202200, Subject 2021ZD0202203 from MOST to MJ; 82101241 and 82360226 grants from NSFC to SW.

## RESOURCE AVAILABILITY

### Lead contact

Further information and requests for resources should be directed to and will be fulfilled by the lead contact.

### Materials availability

This study did not generate new reagents.

### Data and code availability

- All data are available in the main text or the supplementary materials.
- This paper does not report original code.
- Any additional information required to reanalyze the data reported in this paper is available from the lead contact upon request.

## AUTHOR CONTRIBUTIONS

Conceptualization: IAV

Methodology: YL, AB, HZ, PL, YZ, SW, MJ, IAV

Investigation: YL, AB, HZ, PL, QZ, AASS, BW, SSN, XL, DL, SH, RK, WK, YZ, MJ, IAV

Visualization: YL, AB, HZ, PL, DL, IAV

Funding acquisition: SW, MJ, IAV

Project administration: IAV

Supervision: YZ, SW, MJ, IAV

Writing – original draft: YL, MJ, IAV

Writing – review & editing: YL, YZ, SW, MJ, IAV

All authors reviewed and contributed to the manuscript.

## SUPPLEMENTAL INFORMATION

Extended Materials and Methods

Figures S1 to S7

Tables S1 to S3

Movies S1 to S5

## REFERENCES

1 Worldwide trends in underweight and obesity from 1990 to 2022: a pooled analysis of 3663 population-representative studies with 222 million children, adolescents, and adults. Lancet 403, 1027–1050, doi:10.1016/s0140-6736(23)02750-2 (2024).

2 Kusminski, C. M. et al. Transforming obesity: The advancement of multi-receptor drugs. Cell 187, 3829–3853, doi:10.1016/j.cell.2024.06.003 (2024).

3 Chakhtoura, M. et al. Pharmacotherapy of obesity: an update on the available medications and drugs under investigation. eClinicalMedicine 58, doi:10.1016/j.eclinm.2023.101882 (2023).

4 Lenharo, M. Anti-obesity drugs’ side effects: what we know so far. Nature 622, 682, doi:10.1038/d41586-023-03183-3 (2023).

5 Blüher, M. et al. New insights into the treatment of obesity. Diabetes, obesity & metabolism 25, 2058–2072, doi:10.1111/dom.15077 (2023).

6 Waterson, M. J. & Horvath, T. L. Neuronal Regulation of Energy Homeostasis: Beyond the Hypothalamus and Feeding. Cell metabolism 22, 962–970, doi:10.1016/j.cmet.2015.09.026 (2015).

7 Tran, L. T. et al. Hypothalamic control of energy expenditure and thermogenesis. Experimental & molecular medicine 54, 358–369, doi:10.1038/s12276-022-00741-z (2022).

8 Ghamari-Langroudi, M., Srisai, D. & Cone, R. D. Multinodal regulation of the arcuate/paraventricular nucleus circuit by leptin. Proceedings of the National Academy of Sciences of the United States of America 108, 355–360, doi:10.1073/pnas.1016785108 (2011).

9 Modi, M. E. et al. Melanocortin Receptor Agonists Facilitate Oxytocin-Dependent Partner Preference Formation in the Prairie Vole. Neuropsychopharmacology : official publication of the American College of Neuropsychopharmacology 40, 1856–1865, doi:10.1038/npp.2015.35 (2015).

10 Blevins, J. E. & Ho, J. M. Role of oxytocin signaling in the regulation of body weight. Reviews in Endocrine and Metabolic Disorders 14, 311–329, doi:10.1007/s11154-013-9260-x (2013).

11 Liu, H. et al. Transgenic mice expressing green fluorescent protein under the control of the melanocortin-4 receptor promoter. J Neurosci 23, 7143–7154, doi:10.1523/jneurosci.23-18-07143.2003 (2003).

12 McCormack, S. E., Blevins, J. E. & Lawson, E. A. Metabolic Effects of Oxytocin. Endocrine reviews 41, 121–145, doi:10.1210/endrev/bnz012 (2020).

13 Kublaoui, B. M., Gemelli, T., Tolson, K. P., Wang, Y. & Zinn, A. R. Oxytocin Deficiency Mediates Hyperphagic Obesity of Sim1 Haploinsufficient Mice. Molecular Endocrinology 22, 1723–1734, doi:10.1210/me.2008-0067 (2008).

14 Xi, D. et al. Ablation of Oxytocin Neurons Causes a Deficit in Cold Stress Response. Journal of the Endocrine Society 1, 1041–1055, doi:10.1210/js.2017-00136 (2017).

15 Atasoy, D., Betley, J. N., Su, H. H. & Sternson, S. M. Deconstruction of a neural circuit for hunger. Nature 488, 172–177, doi:10.1038/nature11270 (2012).

16 Inada, K., Tsujimoto, K., Yoshida, M., Nishimori, K. & Miyamichi, K. Oxytocin signaling in the posterior hypothalamus prevents hyperphagic obesity in mice. eLife 11, doi:10.7554/eLife.75718 (2022).

17 Gruber, T. et al. High-calorie diets uncouple hypothalamic oxytocin neurons from a gut-to-brain satiation pathway via κ-opioid signaling. Cell reports 42, 113305, doi:10.1016/j.celrep.2023.113305 (2023).

18 Camerino, C. Low sympathetic tone and obese phenotype in oxytocin-deficient mice. Obesity (Silver Spring, Md.) 17, 980–984, doi:10.1038/oby.2009.12 (2009).

19 de Oliveira, M. et al. Pitfalls and challenges of the purinergic signaling cascade in obesity. Biochemical pharmacology 182, 114214, doi:10.1016/j.bcp.2020.114214 (2020).

20 Burnstock, G. Purinergic signalling in endocrine organs. Purinergic signalling 10, 189–231, doi:10.1007/s11302-013-9396-x (2014).

21 Zarrinmayeh, H. & Territo, P. R. Purinergic Receptors of the Central Nervous System: Biology, PET Ligands, and Their Applications. Molecular imaging 19, 1536012120927609, doi:10.1177/1536012120927609 (2020).

22 Huang, Z. et al. From purines to purinergic signalling: molecular functions and human diseases. Signal transduction and targeted therapy 6, 162, doi:10.1038/s41392-021-00553-z (2021).

23 Chen, Y. et al. Spatiotemporally selective astrocytic ATP dynamics encode injury information sensed by microglia following brain injury in mice. Nature neuroscience, doi:10.1038/s41593-024-01680-w (2024).

24 Li, H. et al. Astrocytes release ATP/ADP and glutamate in flashes via vesicular exocytosis. Molecular psychiatry, doi:10.1038/s41380-024-02851-8 (2024).

25 Haynes, S. E. et al. The P2Y12 receptor regulates microglial activation by extracellular nucleotides. Nature neuroscience 9, 1512–1519, doi:10.1038/nn1805 (2006).

26 Cserép, C. et al. Microglia monitor and protect neuronal function through specialized somatic purinergic junctions. Science (New York, N.Y.) 367, 528–537, doi:10.1126/science.aax6752 (2020).

27 Butovsky, O. et al. Identification of a unique TGF-β-dependent molecular and functional signature in microglia. Nat Neurosci 17, 131–143, doi:10.1038/nn.3599 (2014).

28 Zhang, Y. et al. An RNA-sequencing transcriptome and splicing database of glia, neurons, and vascular cells of the cerebral cortex. J Neurosci 34, 11929–11947, doi:10.1523/jneurosci.1860-14.2014 (2014).

29 Lou, N. et al. Purinergic receptor P2RY12-dependent microglial closure of the injured blood-brain barrier. Proc Natl Acad Sci U S A 113, 1074–1079, doi:10.1073/pnas.1520398113 (2016).

30 Hu, L. et al. Platelets Express Activated P2Y(12) Receptor in Patients With Diabetes Mellitus. Circulation 136, 817–833, doi:10.1161/circulationaha.116.026995 (2017).

31 Valdearcos, M. et al. Microglial Inflammatory Signaling Orchestrates the Hypothalamic Immune Response to Dietary Excess and Mediates Obesity Susceptibility. Cell Metab 26, 185–197.e183, doi:10.1016/j.cmet.2017.05.015 (2017).

32 Wang, Z. et al. Saturated fatty acids activate microglia via Toll-like receptor 4/NF-κB signalling. The British journal of nutrition 107, 229–241, doi:10.1017/s0007114511002868 (2012).

33 Zhang, X. et al. Hypothalamic IKKbeta/NF-kappaB and ER stress link overnutrition to energy imbalance and obesity. Cell 135, 61–73, doi:10.1016/j.cell.2008.07.043 (2008).

34 Shih, R. H., Wang, C. Y. & Yang, C. M. NF-kappaB Signaling Pathways in Neurological Inflammation: A Mini Review. Front Mol Neurosci 8, 77, doi:10.3389/fnmol.2015.00077 (2015).

35 Hudson, W. H. et al. Cryptic glucocorticoid receptor-binding sites pervade genomic NF-κB response elements. Nature communications 9, 1337, doi:10.1038/s41467-018-03780-1 (2018).

36 Liu, Y. et al. Functional redundancy between glucocorticoid and mineralocorticoid receptors in mature corticotropin-releasing hormone neurons protects from obesity. Obesity 32, 1885–1896, 10.1002/oby.24116 (2024).

37 Bhuyan, B. K. et al. Tissue distribution of streptozotocin (NSC-85998). Cancer chemotherapy reports 58, 157–165 (1974).

38 Lein, E. S. et al. Genome-wide atlas of gene expression in the adult mouse brain. Nature 445, 168–176, doi:10.1038/nature05453 (2007).

39 Bennett, M. L. et al. New tools for studying microglia in the mouse and human CNS. Proceedings of the National Academy of Sciences of the United States of America 113, E1738–1746, doi:10.1073/pnas.1525528113 (2016).

40 Campbell, J. N. et al. A molecular census of arcuate hypothalamus and median eminence cell types. Nature neuroscience 20, 484–496, doi:10.1038/nn.4495 (2017).

41 Lee, S. D. et al. IDOL regulates systemic energy balance through control of neuronal VLDLR expression. Nature metabolism 1, 1089–1100, doi:10.1038/s42255-019-0127-7 (2019).

42 Rossi, M. A. et al. Obesity remodels activity and transcriptional state of a lateral hypothalamic brake on feeding. Science 364, 1271–1274, doi:10.1126/science.aax1184 (2019).

43 Lei, Y. et al. Region-specific transcriptomic responses to obesity and diabetes in macaque hypothalamus. Cell metabolism 36, 438–453.e436, doi:10.1016/j.cmet.2024.01.003 (2024).

44 Chen, R., Wu, X., Jiang, L. & Zhang, Y. Single-Cell RNA-Seq Reveals Hypothalamic Cell Diversity. Cell reports 18, 3227–3241, doi:10.1016/j.celrep.2017.03.004 (2017).

45 Wen, S. et al. Spatiotemporal single-cell analysis of gene expression in the mouse suprachiasmatic nucleus. Nature neuroscience 23, 456–467, doi:10.1038/s41593-020-0586-x (2020).

46 Hajdarovic, K. H. et al. Single-cell analysis of the aging female mouse hypothalamus. Nature aging 2, 662–678, doi:10.1038/s43587-022-00246-4 (2022).

47 Xu, Y. et al. Glutamate mediates the function of melanocortin receptor 4 on Sim1 neurons in body weight regulation. Cell metabolism 18, 860–870, doi:10.1016/j.cmet.2013.11.003 (2013).

48 Shah, B. P. et al. MC4R-expressing glutamatergic neurons in the paraventricular hypothalamus regulate feeding and are synaptically connected to the parabrachial nucleus. Proceedings of the National Academy of Sciences of the United States of America 111, 13193–13198, doi:10.1073/pnas.1407843111 (2014).

49 Fenselau, H. et al. A rapidly acting glutamatergic ARC → PVH satiety circuit postsynaptically regulated by α-MSH. Nature neuroscience 20, 42–51, doi:10.1038/nn.4442 (2017).

50 Krashes, M. J., Lowell, B. B. & Garfield, A. S. Melanocortin-4 receptor-regulated energy homeostasis. Nature neuroscience 19, 206–219, doi:10.1038/nn.4202 (2016).

51 Sutton, G. M., Duos, B., Patterson, L. M. & Berthoud, H. R. Melanocortinergic modulation of cholecystokinin-induced suppression of feeding through extracellular signal-regulated kinase signaling in rat solitary nucleus. Endocrinology 146, 3739–3747, doi:10.1210/en.2005-0562 (2005).

52 He, S. & Tao, Y. X. Defect in MAPK signaling as a cause for monogenic obesity caused by inactivating mutations in the melanocortin-4 receptor gene. International journal of biological sciences 10, 1128–1137, doi:10.7150/ijbs.10359 (2014).

53 Wang, Y. & Prywes, R. Activation of the c-fos enhancer by the erk MAP kinase pathway through two sequence elements: the c-fos AP-1 and p62TCF sites. Oncogene 19, 1379–1385, doi:10.1038/sj.onc.1203443 (2000).

54 Glas, E., Mückter, H., Gudermann, T. & Breit, A. Exchange factors directly activated by cAMP mediate melanocortin 4 receptor-induced gene expression. Scientific reports 6, 32776, doi:10.1038/srep32776 (2016).

55 Büch, T. R., Heling, D., Damm, E., Gudermann, T. & Breit, A. Pertussis toxin-sensitive signaling of melanocortin-4 receptors in hypothalamic GT1-7 cells defines agouti-related protein as a biased agonist. J Biol Chem 284, 26411–26420, doi:10.1074/jbc.M109.039339 (2009).

56 Mohammad, S. et al. Constitutive traffic of melanocortin-4 receptor in Neuro2A cells and immortalized hypothalamic neurons. The Journal of biological chemistry 282, 4963–4974, doi:10.1074/jbc.M608283200 (2007).

57 Jawien, J. et al. Ticlopidine attenuates progression of atherosclerosis in apolipoprotein E and low density lipoprotein receptor double knockout mice. European journal of pharmacology 556, 129–135, doi:10.1016/j.ejphar.2006.11.028 (2007).

58 Vaughan, K. L. & Mattison, J. A. Obesity and Aging in Humans and Nonhuman Primates: A Mini-Review. Gerontology 62, 611–617, doi:10.1159/000445800 (2016).

59 DeLany, J. P., Hansen, B. C., Bodkin, N. L., Hannah, J. & Bray, G. A. Long-term calorie restriction reduces energy expenditure in aging monkeys. The journals of gerontology. Series A, Biological sciences and medical sciences 54, B5–11; discussion B12-13, doi:10.1093/gerona/54.1.b5 (1999).

60 Cefalu, W. T. et al. Caloric restriction and cardiovascular aging in cynomolgus monkeys (Macaca fascicularis): metabolic, physiologic, and atherosclerotic measures from a 4-year intervention trial. *The journals of gerontology. Series A*, Biological sciences and medical sciences 59, 1007–1014, doi:10.1093/gerona/59.10.b1007 (2004).

61 Colman, R. J. et al. Caloric restriction delays disease onset and mortality in rhesus monkeys. Science (New York, N.Y.) 325, 201–204, doi:10.1126/science.1173635 (2009).

62 Weiss, R. et al. Obesity and the metabolic syndrome in children and adolescents. The New England journal of medicine 350, 2362–2374, doi:10.1056/NEJMoa031049 (2004).

63 Huang, Y. et al. Lipid profiling identifies modifiable signatures of cardiometabolic risk in children and adolescents with obesity. Nat Med 31, 294–305, doi:10.1038/s41591-024-03279-x (2025).

64 Hasan, M. T. et al. A Fear Memory Engram and Its Plasticity in the Hypothalamic Oxytocin System. Neuron 103, 133–146.e138, doi:10.1016/j.neuron.2019.04.029 (2019).

65 Di Virgilio, F., Vultaggio-Poma, V., Falzoni, S. & Giuliani, A. L. Extracellular ATP: A powerful inflammatory mediator in the central nervous system. Neuropharmacology 224, 109333, doi:10.1016/j.neuropharm.2022.109333 (2023).

66 Zhang, W., Xiao, D., Mao, Q. & Xia, H. Role of neuroinflammation in neurodegeneration development. Signal transduction and targeted therapy 8, 267, doi:10.1038/s41392-023-01486-5 (2023).

67 Wei, B. et al. Microglia in the hypothalamic paraventricular nucleus sense hemodynamic disturbance and promote sympathetic excitation in hypertension. Immunity 57, 2030–2042.e2038, doi:10.1016/j.immuni.2024.07.011 (2024).

68 Sivukhina, E. et al. Expression of corticosteroid-binding protein in the human hypothalamus, co-localization with oxytocin and vasopressin. 38, 253–259 (2006).

69 Li, C. et al. Defined Paraventricular Hypothalamic Populations Exhibit Differential Responses to Food Contingent on Caloric State. Cell metabolism 29, 681–694.e685, doi:10.1016/j.cmet.2018.10.016 (2019).

70 Krashes, M. J., Shah, B. P., Koda, S. & Lowell, B. B. Rapid versus delayed stimulation of feeding by the endogenously released AgRP neuron mediators GABA, NPY, and AgRP. Cell metabolism 18, 588–595, doi:10.1016/j.cmet.2013.09.009 (2013).

71 Paul, M. et al. Clopidogrel Administration Impairs Post-Stroke Learning and Memory Recovery in Mice. International journal of molecular sciences 24, doi:10.3390/ijms241411706 (2023).

72 Keller, L. A., Merkel, O. & Popp, A. Intranasal drug delivery: opportunities and toxicologic challenges during drug development. Drug delivery and translational research 12, 735–757, doi:10.1007/s13346-020-00891-5 (2022).

73 Patel, D. & Thakkar, H. Formulation considerations for improving intranasal delivery of CNS acting therapeutics. Therapeutic delivery 13, 371–381, doi:10.4155/tde-2022-0018 (2022).

74 Jeong, S. H., Jang, J. H. & Lee, Y. B. Drug delivery to the brain via the nasal route of administration: exploration of key targets and major consideration factors. Journal of pharmaceutical investigation 53, 119–152, doi:10.1007/s40005-022-00589-5 (2023).

75 Gómez-Pinedo, U. et al. Intranasal Administration of Undifferentiated Oligodendrocyte Lineage Cells as a Potential Approach to Deliver Oligodendrocyte Precursor Cells into Brain. International journal of molecular sciences 22, doi:10.3390/ijms221910738 (2021).

76 Mansour, A., Bachelot-Loza, C., Nesseler, N., Gaussem, P. & Gouin-Thibault, I. P2Y(12) Inhibition beyond Thrombosis: Effects on Inflammation. International journal of molecular sciences 21, doi:10.3390/ijms21041391 (2020).

77 Wijeyeratne, Y. D. & Heptinstall, S. Anti-platelet therapy: ADP receptor antagonists. British journal of clinical pharmacology 72, 647–657, doi:10.1111/j.1365-2125.2011.03999.x (2011).

78 Ferguson, C. S. & Tyndale, R. F. Cytochrome P450 enzymes in the brain: emerging evidence of biological significance. Trends in pharmacological sciences 32, 708–714, doi:10.1016/j.tips.2011.08.005 (2011).

79 Blackmore, K. A., Jeong, J. K. & Young, C. N. A Novel Oxytocin Expressing Microglia Population in the Brain Subfornical Organ. The FASEB Journal 32, 598.599–598.599, 10.1096/fasebj.2018.32.1_supplement.598.9 (2018).

80 Maejima, Y. et al. Identification of oxytocin expression in human and murine microglia. Progress in neuro-psychopharmacology & biological psychiatry 119, 110600, doi:10.1016/j.pnpbp.2022.110600 (2022).

81 Wang, Y. et al. Accurate quantification of astrocyte and neurotransmitter fluorescence dynamics for single-cell and population-level physiology. Nat Neurosci 22, 1936–1944, doi:10.1038/s41593-019-0492-2 (2019).

82 Wu, Z. et al. An obligate role of oxytocin neurons in diet induced energy expenditure. PloS one 7, e45167, doi:10.1371/journal.pone.0045167 (2012).

83 Madisen, L. et al. A robust and high-throughput Cre reporting and characterization system for the whole mouse brain. Nature neuroscience 13, 133–140, doi:10.1038/nn.2467 (2010).

84 Platt, R. J. et al. CRISPR-Cas9 knockin mice for genome editing and cancer modeling. Cell 159, 440–455, doi:10.1016/j.cell.2014.09.014 (2014).

85 Tronche, F. et al. Disruption of the glucocorticoid receptor gene in the nervous system results in reduced anxiety. Nature genetics 23, 99–103, doi:10.1038/12703 (1999).

86 Yu, Y. et al. Interneuron origin and molecular diversity in the human fetal brain. Nature neuroscience 24, 1745–1756, doi:10.1038/s41593-021-00940-3 (2021).

87 Machnicki, A. L. et al. Altered IGF-I activity and accelerated bone elongation in growth plates precede excess weight gain in a mouse model of juvenile obesity. Journal of applied physiology (Bethesda, Md.: 1985) 132, 511–526, doi:10.1152/japplphysiol.00431.2021 (2022).

88 Oraha, J., Enriquez, R. F., Herzog, H. & Lee, N. J. Sex-specific changes in metabolism during the transition from chow to high-fat diet feeding are abolished in response to dieting in C57BL/6J mice. International journal of obesity (2005) 46, 1749–1758, doi:10.1038/s41366-022-01174-4 (2022).

89 Maric, I. et al. Sex and Species Differences in the Development of Diet-Induced Obesity and Metabolic Disturbances in Rodents. Frontiers in nutrition 9, 828522, doi:10.3389/fnut.2022.828522 (2022).

90 Vickers, S. P., Jackson, H. C. & Cheetham, S. C. The utility of animal models to evaluate novel anti-obesity agents. British journal of pharmacology 164, 1248–1262, doi:10.1111/j.1476-5381.2011.01245.x (2011).

91 Giovannelli, L., Shiromani, P. J., Jirikowski, G. F. & Bloom, F. E. Oxytocin neurons in the rat hypothalamus exhibit c-fos immunoreactivity upon osmotic stress. Brain research 531, 299–303, doi:10.1016/0006-8993(90)90789-e (1990).

