## Supplementary Information for "Remodeling of purinergic signaling in the paraventricular hypothalamus promotes hyperphagic obesity and insulin resistance"

**This PDF file includes:**

Extended Materials and Methods
Figures S1 to S7
Tables S1 to S3

**Other Supplemental Information for this manuscript includes the following:**

Movies S1 to S5

### EXTENDED MATERIALS AND METHODS

#### Cell Culture

The mouse hypothalamic GT1-7 (chosen because they express MC4R [55]) and human kidney HEK293T cell lines were purchased from the Cell Bank of Shanghai Institutes for Biological Sciences, China and cultured in DMEM (Gibco, 11960044) containing 10% fetal bovine serum (Hyclone, SV30087.03), 100 U/ml penicillin and 100 U/ml streptomycin at 37 °C and 5% $CO_2$. The growth of cells was observed daily, with the culture medium replaced daily or every other day according to the conditions of the cells.

$GRkd_1$ and $GRkd_2$ vectors were constructed by subcloning shRNAs targeting *Nr3c1* (UUU GCU CCU GAU CUG AUU AUU and UGG AUA AGU CCA UGA GUA UUG, respectively) into pSLenti-U6-shRNA-CMV-EGFP-F2A-Puro-WPRE plasmid. For *in vitro* $P2Y_{12}$ overexpression, mouse *P2ry12* cDNA was subcloned into pCDH-EF1-copGFP-T2A-Puro (Addgene, 72263, a gift from Kazuhiro Oka) at EcoRI and BamHI sites. The resulting Lenti-$P2ry_{12}$OE, $GRkd_1$ and $GRkd_2$ constructs were packaged, transduced in GT1-7 cells and 72 h later FACS-sorted, followed by verification of single clones by qRT-PCR. GT1-7 cells were treated for 1 h by vehicle (0.001% DMSO) or 100 nM α-MSH (APExBio, A1025) to induce cAMP and downstream targets [56]. For cAMP ELISA analysis, prior to α-MSH addition, cells were pretreated by 10 μM $P2Y_{12}$ inhibitor 2-methylthio-AMP (2MeSAMP, MCE, HY-125989) or vehicle (0.1% DMSO) for 12 hours.

Two guide RNAs (gRNAs) targeting *P2ry12* with highest scores and lowest putative off-target sites (ACC GCT ACC TGA AGA CCA CC and CGT TCA GTG ACG TCA GCC AT) were predicted using http://chopchop.cbu.uib.no and validated (LY, AB, SSN, DL, SH, YZ, IAV in preparation).

#### Animal models

Mice were maintained on a 12/12 hours light/dark cycle with free access to water and chow (11.1% kcal from fat, Jiangsu Xietong Pharmaceutical Bio-engineering Co, 1010088) or HFD food (60% kcal from fat, Jiangsu Xietong Pharmaceutical Bio-engineering Co, XTHF60-1 and XTHF60 for the experiments with organotypic slices and all others, respectively), with the same batch per experiment. The diet induced obesity (DIO) mouse models were fed the HFD for periods indicated in the figure legends, starting from Week 5, unless otherwise stated. Blinding was applied prior to and during the group allocation and the course of the experiments, as well as during assessment and analysis of outcomes, unless, for example, due to strong metabolic phenotypes, researchers could not be blinded in regard to experimental groups Wherever possible, RAND() function in Excel was used for randomization. To reduce potential variability, laboratory animals were housed in standard specific pathogen free (SPF) grade animal rooms. Death of animals during the experiments was used as an exclusion criterion. While experiments with knock-out of GR, over-expression of $P2Y_{12}$, initial oral and nasal administration of inhibitors were conducted on mice of both sexes (see legends for details), due to the faster and more robust HFD-mediated induction of weight gain in males, they were used in all other experiments. In particular, predominantly used male mice in HFD-induced obesity studies and weight-loss upon drug administration due to several factors. Firstly, male mice, particularly the C57BL/6J strain, are more prone to obesity, hyperinsulinemia, hyperglycemia, and hypertension when fed with a HFD, in contrast to females exhibiting resistance to these effects, thus making males a more consistent model for studying obesity-related outcomes [88].

Secondly, male rodents tend to consume more food and gain weight more rapidly on high-fat diets compared to females, with the latter displaying delayed weight gain and fewer metabolic complications, attributed to higher energy expenditure and lower hyperphagia [89]. Finally, acute studies often use males to quickly assess the potency, efficacy, and potential side effects of compounds [90]. Our initial nasal administration experiments revealed effect sizes reaching 1.46, 1.19, 1.58 and 1.45 for prasugrel, ticlopidine, ticagrelor and cangrelor, respectively. This allowed us to optimize our study design by reducing sample sizes to 5-6 per group for subsequent experiments, while maintaining the statistical rigor and ensuring ethical and efficient use of animals, in agreement with previously reported effect sizes for body mass and other parameters in mice subjected to HFD with sample sizes as low as 5 per group [87]. Moreover, since nasal application of $P2Y_{12}$ inhibitors was evaluated *in vivo* for the first time in the present study, the initial intention was to demonstrate the effect of inhibition of the purinergic pathway by various reversible or irreversible compounds and gather basic evidence regarding their use at different concentrations. Metabolic profiles were analyzed in Comprehensive Lab Animal Monitoring System (CLAMS, Columbus Instruments). Fat/lean mass composition was measured by magnetic resonance imaging (Echo-MRI) system MesoMR23-060H–I (NIUMAG) according to the manufacturer's protocol at the Instrumental Analysis Center of Shanghai Jiao Tong University. For glucose (GTT) and insulin tolerance tests (ITT), mice were fasted for 16 h and 4 h followed by intraperitoneal injections of 2 g/kg D-glucose (Sigma-Aldrich, GT528) and 0.5 U/kg insulin (Sigma-Aldrich, I9278), respectively. Blood glucose levels were measured at time points 0, 15, 30, 60, 90 and 120 min using the ACCU-CHEK glucometer (Roche). For the diabetic model, single intraperitoneal injection of vehicle or 200 mg/kg streptozotocin in 0.1 M citrate buffer were administered to male mice followed by blood glucose level measurements, with the threshold of > 300 mg/dl (16.7 mmol/l) considered as diabetic. This diabetic model with systemic administration had been chosen to avoid direct cytotoxicity on the PVH, because streptozotocin does not pass through the blood-brain barrier [37=Bhuyan 1974].

Cynomolgus monkeys (*Macaca fascicularis*) were fed with a diet consisting of monkey chow, supplemented by seasonal fresh fruits and vegetables. Body Condition Score (BCS) was used as a readout of voracious appetite and obesity onset, with the values above 4.5 considered as extremely obese for this species. Hence, when the BCS of *M. fascicularis* became equal or greater than 4, the diet was meticulously regulated by a veterinarian with a total daily amount of 410 g including chow food (Xietong Shenwu, Q/321081 YZXT 01-2023) in the morning (10:00-10:30) and in the afternoon (16:00-16:30), fruits and vegetables at 11:00-11:30 and peanuts at 17:00. Prior to the short-term experiment, the six animals were maintained under these calorie restriction conditions receiving maximum 150 g of lab monkey maintenance chow food in the morning supplemented by 150 g of fruits and vegetables, which was followed by 100 g monkey chow in the afternoon supplemented by 10 g peanuts. During the experiment, we increased the afternoon chow portion to 200 g. Prior to the start of the long-term experiment, the three animals received 150 g of chow food in the morning supplemented by 100 g of fruits and vegetables, which was followed by 150 g monkey chow in the afternoon supplemented by 10 g peanuts. During the experiment, morning chow food amount was increased to 200 g, fruits and vegetables—to 120 g and afternoon chow food—to 250 g. The age, initial weights and BCS indices for each animal are indicated in **Table S2**. Fasting blood was sampled before the first meal, firstly on the day prior to the termination of calorie restriction and secondly on the $157^{th}$, last day of cangrelor treatment, followed by analysis on Beckman Coulter AU5800 with kits #AUZ3139, AUZ3204, AUZ3016, AUZ3605, KP629, HY5148, 2569, 2629, 2024091401, 2024050701, 40515B11, AE5196, R10241001, AUZ2732, AUZ3370, AUZ3583, AUZ2974, AUZ3315, AUZ3365, AUZ3724, AUZ3223, AUZ3325 and AUZ3756 for total cholesterol, high- and low-density lipoproteins (HDL and LDL, respectively), glucose, triglycerides, homocysteine, apolipoproteins ApoA1, ApoB, ApoA2, ApoC2, ApoC3, ApoE, lipoprotein-associated phospholipase A2 (LP-PLA2), phosphate, calcium, total or direct bilirubin, total protein, alanine aminotransferase (ALT), aspartate aminotransferase (AST), γ-Glutamyl transferase (GGT), cholinesterase and urea, respectively; Roche Cobas 8000 with kits #78201501, 76420201 and 79975801 for insulin, C-peptide and lipoprotein-associated phospholipase A2NT-proBNP (NT-proBNP), respectively. The normal human reference ranges were indicated according to the hygienic standard WS/T 402-2024 of the National Health Commission of China. No animal in control or inhibitor groups revealed any bleeding episodes (judging by skin and mucosa examinations, as well as by lack of hemoglobin in the stool), abnormalities, or other adverse phenotypes.

**Stereotaxic injections**

Stereotaxic apparatus (RWD) was used to bilaterally deliver 0.5 μL recombinant adeno-associated viral vectors (rAAVs) to the PVH of adult $OXT^{IRES\text{-}Cre}$:Cas9-GFP or $OXT^{IRES\text{-}Cre}$ mice (coordinates relative to bregma: A/P, − 0.85 mm; M/L, ± 0.3 mm; D/V, − 4.75 mm) to knock-out or overexpress $P2Y_{12}$ specifically in $PVH^{OXT}$ neurons, respectively. Briefly, mice were anesthetized by avertin (300 mg/kg) and their heads were fixed in the stereotaxic apparatus. $PVH^{OXT}$:$P2Y_{12}$cko or Scrambled mice received AAV- $gRNA_{P2ry12}$ ($8.16 \times 10^{13}$ vg/ml) or AAV- Scrambled vector ($3.97 \times 10^{13}$ vg/ml), respectively. $PVH^{OXT}$:$P2Y_{12}$ and control mice received AAV-CMV-FLEX-P2ry12 ($4.69 \times 10^{13}$ vector genomes/ml [vg/ml]) and AAV-CMV-FLEX ($3.91 \times 10^{13}$ vg/ml), respectively. All above-mentioned vectors have been constructed and packaged into AAV.CAP-B10 capsids (Vigene Biosciences) to specifically target neurons. Each knock-out vector was designed to contain a cassette with two gRNAs targeting a gene of interest, as described elsewhere [36]. For visualization of Inflares, 0.3 μl AAV2/9- GfaABC1D-ATP1.0 vectors ($1.8 \times 10^{13}$ vg/ml) were injected to the same coordinates of wildtype mice 2 weeks before preparation of organotypic slices. For knocking out *Panx1* in astrocytes, one side of PVH of the *Panx1*$^{fl/fl}$ mice (GemPharmatech, Strain NO. T008074) was injected with AAV-GfaABC1D-Cre ($2.5 \times 10^{13}$ vg/ml), with the contralateral side receiving AAV-GfaABC1D-mCherry ($3.1 \times 10^{13}$ vg/ml).

**Acute organotypic brain slice experiments**

For preparing acute brain slices, mice were anesthetized with avertin two weeks after transduction and were transcardially perfused with 5 ml pre-cooled slicing buffer containing 110 mM choline chloride, 2.5 mM KCl, 1.25 mM $NaH_2PO_4$, 25 mM $NaHCO_3$, 25 mM D-glucose, 7 mM $MgCl_2$ and 0.5 mM $CaCl_2$. The mice were then decapitated, brains were immediately removed, placed in pre-cooled oxygenated slicing buffer oxygenated with 95% $O_2$ + 5% $CO_2$ and sectioned into 300 μm slices using VT1200 vibratome (Leica). The slices were immediately transferred to oxygenated artificial cerebro-spinal fluid (ACSF) containing 125 mM NaCl, 2.5 mM KCl, 1.25 mM $NaH_2PO_4$, 25 mM $NaHCO_3$, 25 mM D-glucose, 1.3 mM $MgCl_2$ and 2 mM $CaCl_2$, recovered at 33°C for at least 30 min before being transferred to a chamber for imaging.

Imaging of acute brain slices at indicated time points after streptozotocin/vehicle injections or HFD/chow diet feeding was conducted using a two-photon laser-scanning microscope (FVMPE-RS; Olympus). Images were acquired using a 25× water immersion objective (XLPLN25XWMP2, 1.05 NA, Olympus), with the field of view set at 500 μm × 500 μm, and imaging frequency set at 10 Hz. For mechanistic studies, 5 mM pannexin blocker probenecid (Invitrogen, P36400) or control vehicle (ACSF) was bath-applied in the chamber for 5 minutes before imaging. Inflare regions of interest (ROIs) in acute slices were identified using AQuA software [81] in MATLAB (MathWorks) and manually validated. The total ROIs within the indicated time period were plotted and merged with the fluorescence image of ATP1.0 using Fiji (ImageJ).

To introduce focal laser injury (FLA), the 920-nm two-photon laser was set at its maximum power and applied to the center of the imaging region (50 μm in diameter, 100% laser power) for 1 s using the Tornado scanning mode. Successful FLA could be observed by the formation of a high fluorescence circle around the FLA region.

**Histological analyses**

The human brain tissue samples were generously donated by the Fudan branch of National Health and Disease Human Brain Tissue Resource Center, the Body Donation Station at Fudan University (Shanghai Red Cross Society). The human postmortem hypothalamus, mouse perigonadal white adipose tissue (WAT) were fixed in 4% paraformaldehyde (PFA), embedded in paraffin, sectioned at a thickness of 5 μm and stained with hematoxylin and eosin (Beyotime, C0105S) according to manufacturer's instructions. In addition, human brain sections were stained by tyramide signal amplification (TSA) kit (WASci, WAS15021) according to manufacturer's instructions at BioMed World using rabbit anti-P2ry12 (Sigma-Aldrich, HPA014518, 1:500), rabbit anti-oxytocin (ImmunoStar, 20068, 1:500), rabbit anti-IBA1 (Cell Signaling Technology, 12943S, 1:2000), rabbit anti-NeuN (Proteintech, 10904-1-AP, 1:500). Eclipse Ci-L (Nikon) and Fluorescence Scanning System Pannoramic MIDI II (3DHISTECH) were used to acquire the images. In detail, paraffin sections were three times incubated in dry oven at 60°C for 1 h, then immersed in xylene for 15 min for deparaffinization. Rehydration was performed by sequentially 5-min-immersing slides in 100% ethanol, 95% ethanol, and 70% ethanol. Next, paraffin sections were three times immersed to tap water for 1 min, then microwaved at high power for 5 min followed by an additional 10 minutes in Tris-EDTA buffer (pH 9.0) at low power for antigen retrieval. After cooling to room temperature (RT), slides were immersed with peroxidase-blocking solution for 5 min to block endogenous peroxidase activity and sealed with the 3% BSA for 30 min. The slides were incubated with the primary antibodies at 4°C overnight, washed thrice in 1× PBS buffer and incubated with horseradish peroxidase (HRP)-conjugated secondary antibodies for 50 min at RT. A quick wash in 1× PBS buffer was followed by incubation with an appropriate fluorophore-TSA conjugate for 10 min at RT. Further, the slides were exposed to microware treatment to strip the tissue-bound primary/secondary antibody complexes to prepare them for labelling by the next antibodies until all markers were labelled. Finally, DAPI (WASci, WAS1301) was applied to the slides in the dark for 10 min at RT followed by mounting with anti-fade mounting medium. The adipose tissue images were acquired by optical microscope (Olympus) followed by adipocyte area analysis by ImageJ and AdipoCount software.

4% PFA-perfused mouse brains were fixed in 4% PFA at 4°C for 4 hours and then dehydrated using a gradient solution of 20% and then 30% sucrose at 4°C overnight, embedded in Tissue-Tek™ optimum cutting temperature compound (Sakura, 4853) and sliced to 10 µm sections using a cryostat (Leica, CM1950). After blocking with 5% serum in 0.3% Triton X-100/PBS, the sections were incubated overnight at 4°C with primary antibodies: rabbit anti-P2ry12 (Sigma-Aldrich, HPA014518, 1:500) or rabbit anti-c-Fos (Abcam, ab222699, 1:200). The following day, the sections were incubated for 1 h at room temperature with 1:500 secondary antibodies conjugated with Dylight-Fluor 405 (Beyotime, A0609) Fluor-647 (Elabscience, E-AB-1075). Images were acquired using a Ni-E A1 HD25 confocal microscope (Nikon).

**FACS**

All procedures were performed on ice, unless otherwise stated. For the PVH microdissection, coronal brain sections were collected using a 0.5 mm mouse stainless steel brain matrix (RWD) slicing immediately after the optical chiasm and 2 mm rostrally from the first cut. Under microscope, the PVH was microdissected from the resulting slice by two scalpels. For tissue digestion, the PVH sample was washed three times with 1× PBS and transferred to 0.5 ml digestion medium containing 2 mg/ml of collagenase IV (Gibco, 17104-019) in Gibco™ Hibernate™-E Medium (A1247601) in a V shape-bottom 2 ml tube. Next, the tissue was gently separated into small pieces by pressing it against the V-shaped bottom or against a lid of the tube using a plunger of a 1 mL syringe followed by gentle mixing using a 1 ml plastic Pasteur pipette. After discarding the supernatant by centrifugation at 150 g for 3 min, the pellet was incubated at 37 °C for 20 min in the Gibco™ Hibernate™-E Medium with 1 mg/ml papain (Shanghai Yuanye Bio-Technology, S10011). 1 ml of Gibco™ Hibernate™-E Medium was added to stop digestion. The released cells were filtered by a 40-µm cell strainer, collected by centrifugation at 500 g for 5 min, re-suspended in 1.5 ml 1× PBS and kept on ice to immediately proceed with FACS sorting (BD FACS Aria III) according to the manufacturer's protocol. In each of the $OXT^{IRES\text{-}Cre}$tdTomato mice, 50 $tdTomato^{+}$ cells were sorted into a tube with lysis buffer from CellAmp™ whole transcriptome amplification kit (Takara, 3734) according to the manufacturer’s protocol, followed by reverse-transcription and subsequent quantitative real-time reverse-transcription PCR (qRT-PCR) on the template of the cDNA.

**Quantitative real-time reverse-transcription PCR**

Total RNA was extracted using TRIzol reagent (Thermo Fisher Scientific, 15596026), reverse-transcribed to cDNA using a reverse transcription reagent kit (Takara, RR047A). qRT-PCR was performed using Hieff® qPCR SYBR Green Master Mix (Yeasen, 11201ES08) on a

Real-Time PCR System (Biorad) using the following primer sequences (forward/reverse): *Fos*, GGG ACA GCC TTT CCT ACT / GAT CTG CGC AAA AGT CCT; *P2ry12*, CCC TGT GCG TCA GAG ACT AC / CAA GCT GTT CGT GAT GAG CC; *Nr3c1*, TGA AGC TTC GGG ATG CCA TT / TTC GAT AGC GGC ATG CTG G; *Nfkb1*, GGA GGC ATG TTC GGT AGT GG / CCC TGC GTT GGA TTT CGT G, *Gapdh*(m), AGG TCG GTG TGA ACG GAT TTG / TGT AGA CCA TGT AGT TGA GGT CA. The $2^{-\Delta Ct}$ method was used to determine relative mRNA levels.

**ELISA**

The adherent cells were washed gently with cold PBS, digested with trypsin, centrifuged at 1,000 g for 5 min, washed 3 times with cold PBS, suspended in cold PBS with protease inhibitors (MCE, HY-K0013) and lysed by repeated freeze-thaw cycles. The supernatant after centrifugation at 2-8°C, 1,500 g for 10 min was analyzed by cyclic adenosine monophosphate (cAMP) ELISA Kit (Elabscience, E-EL-0056) according to the manufacturer's protocol. Origin V9.1 software was used to determine the concentrations based on the generated standard curve.

**Figure S1**

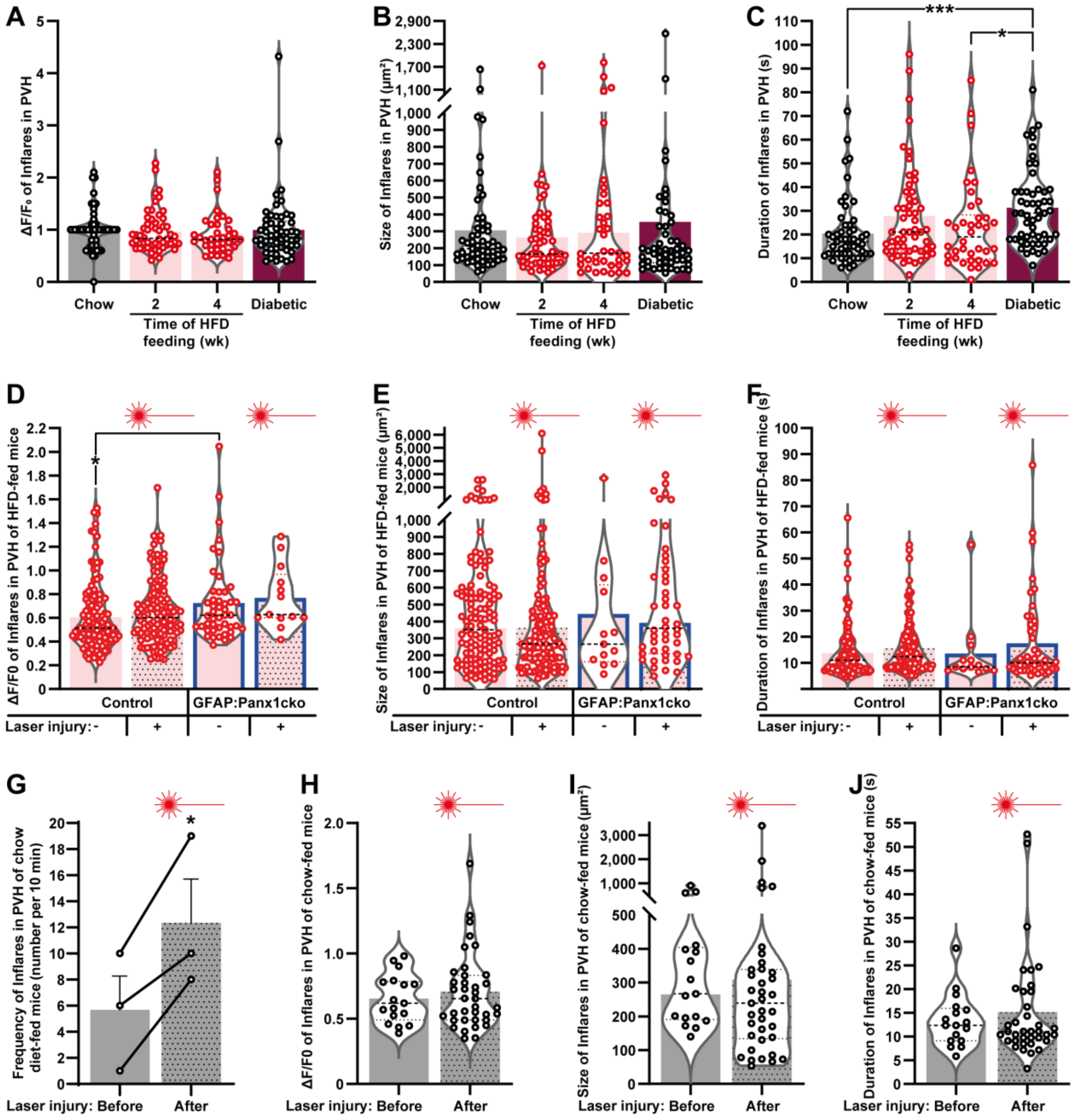


**Figure S1. Assessment of the properties and mechanisms of ATP events in the PVH, related to Figure 1.** (**A-C**) Amplitude ($\Delta F/F_0$, **A**), size (**B**) and duration (**C**) of Inflares in the PVH regions-containing acute hypothalamic slices from male mice fed with chow diet (used as controls for all time points in **Figure 1C**), HFD for 2 and 4 wk, or injected with streptozotocin (Diabetic) (n = 53 Inflares from 9 mice, 54 from 3 mice, 38 from 3 mice and 53 from 3 mice, respectively). (**D-F**) Amplitude (**D**), size (**E**) and duration (**F**) of Inflares before and after focal laser injury in the PVH regions-containing acute hypothalamic slices from male *Panx1*$^{fl/fl}$ mice, with one side of PVH injected with rAAV for astrocytic Cre expression (GFAP:Panx1cko) and the other side with control rAAV. Mice (n = 3) were subjected to HFD-feeding for 2 wk before imaging (n = 119 and 119 Inflares in the control side before and after laser injury, respectively; 46 and 13 Inflares in the GFAP:Panx1cko side before and after laser injury, respectively). For analyses before laser injury 8 slices in total were used; after laser irradiation 4 and 5 slices were used for GFAP:Panx1cko and control side analyses, respectively). (**G-J**) Frequency (**G**, n = 3 slices), amplitude (**H**), size (**I**) and duration (**J**) of laser injury-evoked Inflares in chow diet fed animal-isolated acute PVH slices (n = 17 and 37

Inflares from 3 slices before and after injury, respectively). Error bars represent standard error of means (SEM). *, $p < 0.05$; ***, $p < 0.001$, as analyzed by Kruskal-Wallis test followed by Dunn's multiple comparisons test (**C**,**D**) or paired two-tailed Student’s t-test (**G**).

**Figure S2**

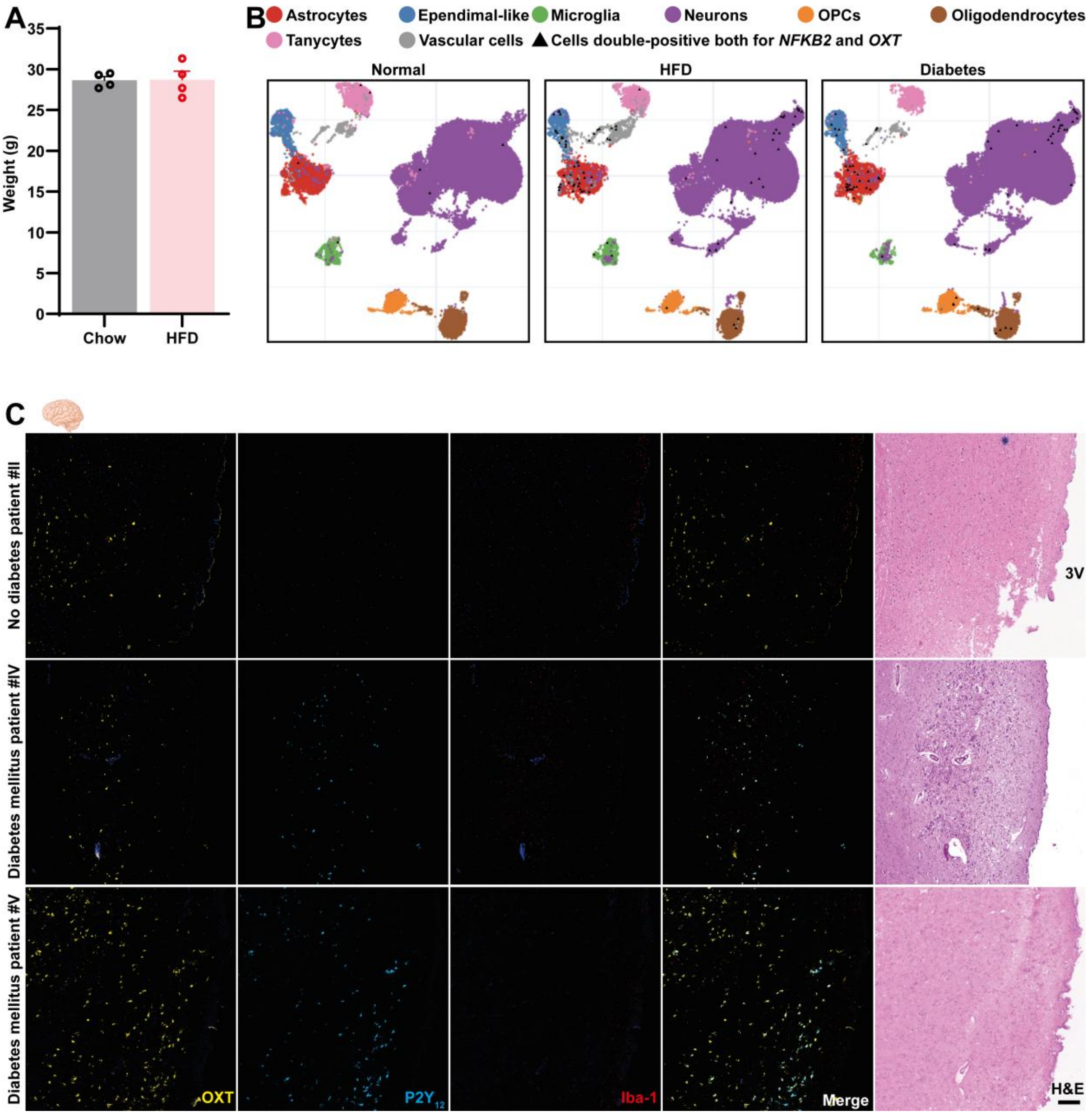


**Figure S2**. **Assessment of purinergic signaling properties within the hypothalamus, related to Figure 2.** (**A**) Weight in chow and HFD-fed female mice from the experiment depicted in **Figure 2B** at the time of analysis of *P2ry12* expression in FACS-isolated $PVH^{OXT}$ neurons (n = 4). (**B**) Co-localization of *NFKB2* with *OXT* (black triangles) in the indicated hypothalamic cellular populations of normal, obese and diabetic rhesus macaques (*Macaca mulatta*), assessed by https://db.cngb.org/cdcp/hca/. OPCs, oligodendrocyte precursor cells. Note that the cellular clusters in the cell maps are not completely homogenic, but contain many inclusions outlined with a different color, i.e. a small number of cells of different populations within the predominant ones, which, however, cluster together. (**C**) Co-localization microphotographs of the PVH-containing *ex vivo* slices from patients with or without diabetes mellitus (n = 3 and 2, respectively). Error bars represent standard error of means (SEM). Scale bar: 200 μm.

**Figure S3**

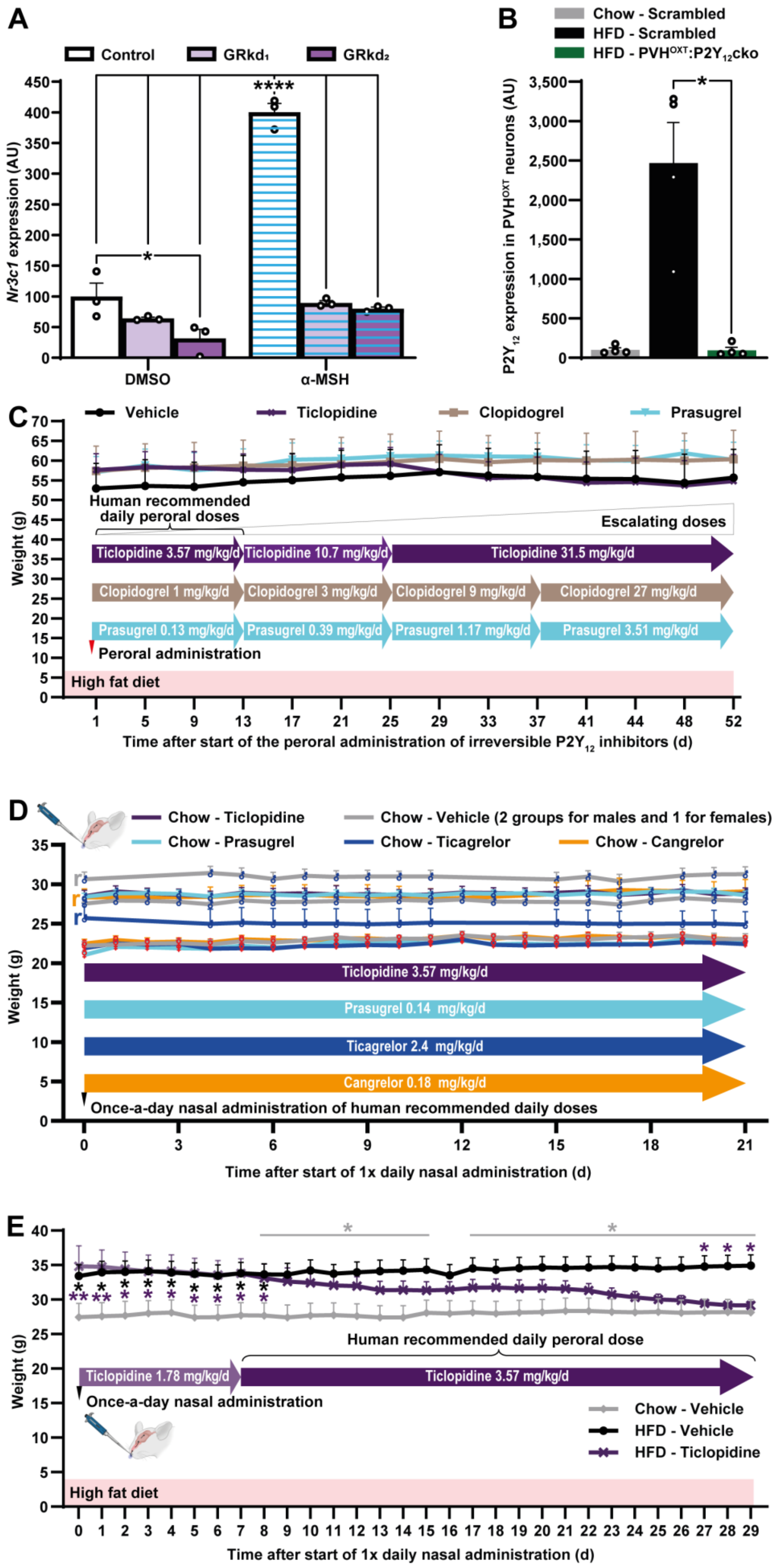

A
Control
GRkd1
GRkd2
Nr3c1 expression (AU)
DMSO
α-MSH
****
*
B
Chow - Scrambled
HFD - Scrambled
HFD - PVHOXT:P2Y12cko
P2Y12 expression in PVHOXT neurons (AU)
*
C
Vehicle
Ticlopidine
Clopidogrel
Prasugrel
Weight (g)
Human recommended daily peroral doses
Escalating doses
Ticlopidine 3.57 mg/kg/d
Ticlopidine 10.7 mg/kg/d
Ticlopidine 31.5 mg/kg/d
Clopidogrel 1 mg/kg/d
Clopidogrel 3 mg/kg/d
Clopidogrel 9 mg/kg/d
Clopidogrel 27 mg/kg/d
Prasugrel 0.13 mg/kg/d
Prasugrel 0.39 mg/kg/d
Prasugrel 1.17 mg/kg/d
Prasugrel 3.51 mg/kg/d
Peroral administration
High fat diet
Time after start of the peroral administration of irreversible P2Y12 inhibitors (d)
D
Chow - Ticlopidine
Chow - Vehicle (2 groups for males and 1 for females)
Chow - Prasugrel
Chow - Ticagrelor
Chow - Cangrelor
Weight (g)
Ticlopidine 3.57 mg/kg/d
Prasugrel 0.14 mg/kg/d
Ticagrelor 2.4 mg/kg/d
Cangrelor 0.18 mg/kg/d
Once-a-day nasal administration of human recommended daily doses
Time after start of 1x daily nasal administration (d)
E
Weight (g)
Human recommended daily peroral dose
Ticlopidine 1.78 mg/kg/d
Ticlopidine 3.57 mg/kg/d
Once-a-day nasal administration
Chow - Vehicle
HFD - Vehicle
HFD - Ticlopidine
High fat diet
Time after start of 1x daily nasal administration (d)

**Figure S3. Loss of glucocorticoid receptors in PVH$^{OXT}$ neurons stimulates P2Y$_{12}$ expression and leads to obesity in mice. Effects of P2Y$_{12}$ receptor deficiency *in vivo* and *in vitro*. Daily resolution of weight dynamics in mice nasally administered with P2Y$_{12}$ inhibitors, related to Figures 3 and 4.** (**A**) qRT-PCR analysis of *Nr3c1* gene expression in GT1-7 hypothalamic cells transduced with lentiviral vectors expressing *Nr3c1* small hairpin RNA (shRNA) with or without 1 h-stimulation by 100 nM α-MSH (n=3). (**B**) Knock-out efficiency quantification in the PVH of PVH$^{OXT}$:P2Y$_{12}$cko male mice (n = 4). (**C**) An experiment to identify the lowest effective peroral doses of prodrugs yielding irreversible inhibitors of P2Y$_{12}$. Body weight in diet-induced obese male mice after peroral administration of vehicle, ticlopidine, clopidogrel, and prasugrel in drinking water at indicated concentrations starting from the human recommended daily peroral doses and gradually increasing 3× after every 3 wk (n = 5, 5, 6, 5, respectively). (**D**) Weight in three separate experiments with once-a-day nasal administration of vehicle or recommended daily doses of P2Y$_{12}$ inhibitors to chow diet-fed wild-type mice. Female groups (n=6) are outlined with respective symbol, reversible inhibitor male groups together with the vehicle controls (n=6) are labelled by "r" before the first time point, n = 10 for irreversible inhibitor and vehicle control male groups. (**E**) Body weight in chow or HFD-fed male mice after once-a-day nasal administration of vehicle or 1.78 mg/kg/d ticlopidine for 1 wk followed by human daily peroral dose 3.57 mg/kg (n = 7, 8, 8, respectively). Error bars represent SEM. *, $p < 0.05$; **, $p < 0.01$; ****, $p<0.0001$ as analyzed by one- (**A**) and two-way (**E**) ANOVA with Holm-Šídák's test or Kruskal-Wallis test followed by Dunn's multiple comparisons test (**B**).

**Figure S4**

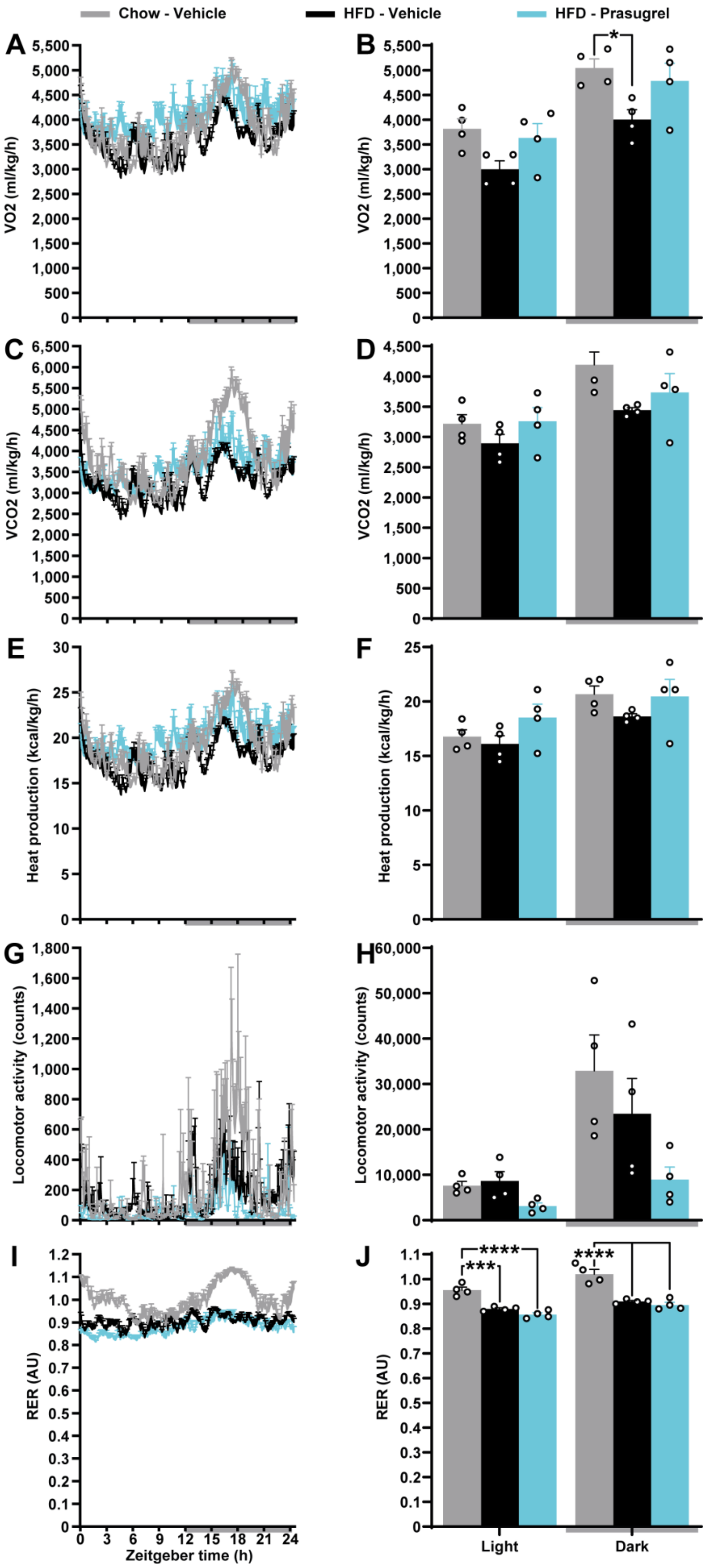

Chow - Vehicle
HFD - Vehicle
HFD - Prasugrel
A
B
C
D
E
F
G
H
I
J
VO2 (ml/kg/h)
VCO2 (ml/kg/h)
Heat production (kcal/kg/h)
Locomotor activity (counts)
RER (AU)
Zeitgeber time (h)
0 3 6 9 12 15 18 21 24
Light
Dark
*
***
****
****

**Figure S4**. **Metabolic analysis in diet-induced obese mice nasally administered with prasugrel, related to Figure 4.** (**A-H**) Raw data plot (**A**,**C**,**E**,**G**,**I**) and 12-hr period metabolic cage analysis (**B**,**D**,**F**,**H**,**I**) of VO2 (**A**,**B**), VCO2 (**C**,**D**), heat production (**E**,**F**), locomotor activity (**G**,**H**) and respiratory exchange rate (RER, **I**,**J**) in chow or HFD male mice nasally administered once per day with vehicle or daily recommended human prasugrel peroral dose of 0.14 mg/kg/d for 30 days (n = 4). Error bars represent SEM. *, $p < 0.05$; ***, $p < 0.001$; ****, $p < 0.0001$ as analyzed by one-way ANOVA with post-hoc Holm-Šídák's test.

**Figure S5**

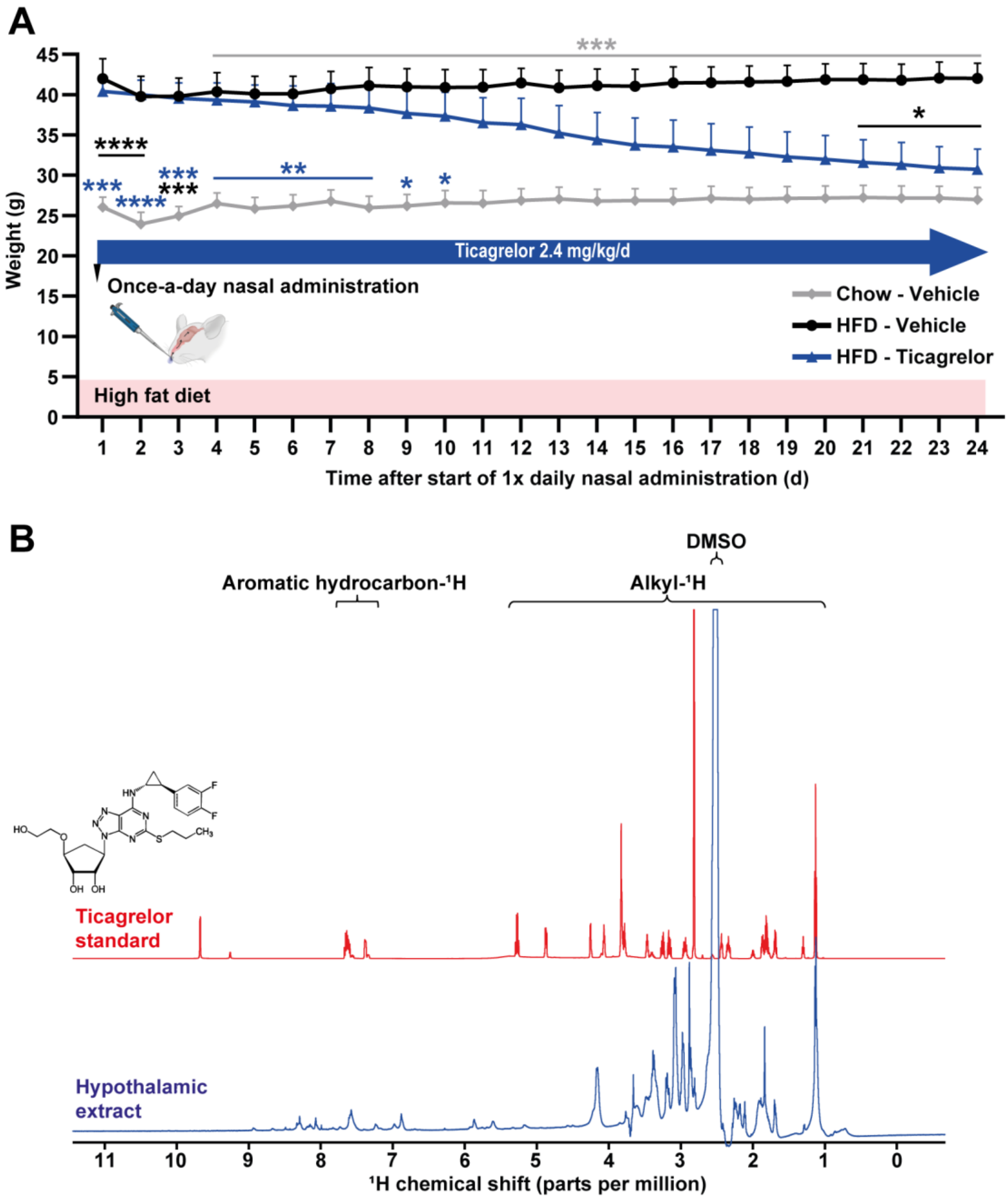


**Figure S5. Abundance in the hypothalamus and anti-obesity activity of nasally administered ticagrelor, related to Figure 5.** (**A**) Daily resolution of weight records in chow and HFD-fed male mice nasally administered once per day with vehicle or daily recommended human ticagrelor dose of 2.4 mg/kg (n = 5). (**B**) After 24 days of daily nasal administration of 2.4 mg/kg ticagrelor, hypothalamic tissues from five male mice were homogenized in deuterated DMSO and mixed, followed by 700 MHz nuclear magnetic resonance (NMR) spectrometer analysis revealing limited shifts in alkyl group peaks, but well-preserved aromatic hydrocarbon peaks characteristic for ticagrelor. Ticagrelor in deuterated DMSO was used as a standard. The water peak has been suppressed. Error bars represent SEM. *, $p < 0.05$; **, $p < 0.01$; ***, $p<0.001$; ****, $p<0.0001$ as analyzed by two-way ANOVA with post-hoc Holm-Šídák's test.

**Figure S6**

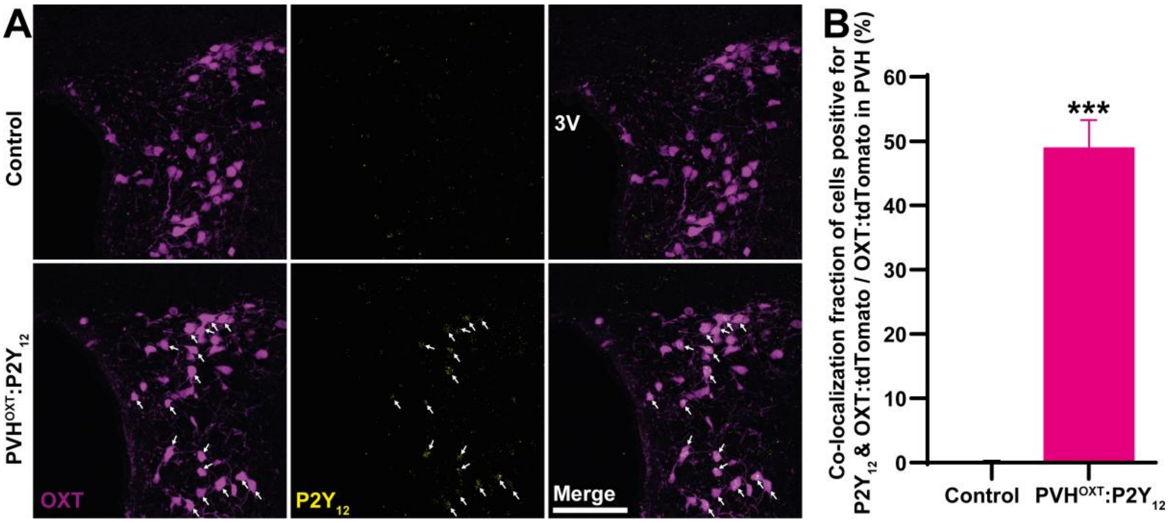


**Figure S6. Expression of P2Y$_{12}$ in PVH$^{OXT}$ neurons of PVH$^{OXT}$:P2Y$_{12}$ mice, related to Figure 7.** (**A,B**) Representative microphotographs (**A**) and quantification (**B**) of P2Y$_{12}$ immunofluorescence in the PVH of PVH$^{OXT}$:P2Y$_{12}$ male mice. Error bars represent SEM. ***, $p < 0.001$ as assessed by unpaired 2-tailed Student's t-test. Scale bar: 100 µm.

**Figure S7**

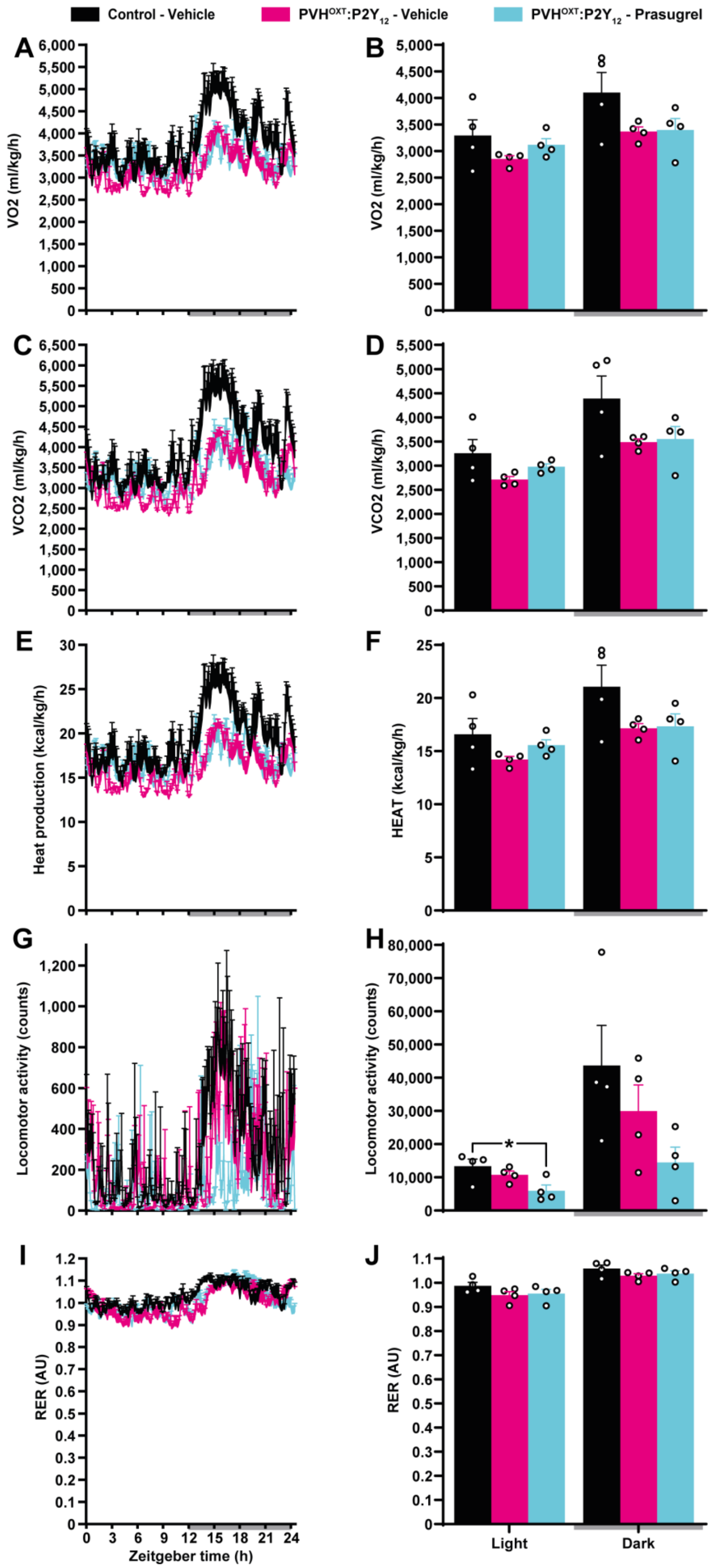

Control - Vehicle
PVH$^{OXT}$:P2Y$_{12}$ - Vehicle
PVH$^{OXT}$:P2Y$_{12}$ - Prasugrel
A
B
VO2 (ml/kg/h)
C
D
VCO2 (ml/kg/h)
E
F
Heat production (kcal/kg/h)
HEAT (kcal/kg/h)
G
H
Locomotor activity (counts)
I
J
RER (AU)
Zeitgeber time (h)
Light
Dark

**Figure S7. Metabolic analysis in PVH$^{OXT}$:P2Y$_{12}$ mice nasally administered with prasugrel, related to Figures 7.** (**A-H**) Raw data plot (**A**,**C**,**E**,**G**,**I**) and 12-h period metabolic cage analysis (**B**,**D**,**F**,**H**,**I**) of VO2 (**A**,**B**), VCO2 (**C**,**D**), heat production (**E**,**F**), locomotor activity (**G**,**H**) and RER (**I**,**J**) in chow diet-fed PVH$^{OXT}$:P2Y$_{12}$ male mice nasally administered once per day with vehicle or daily recommended human prasugrel dose of 0.14 mg/kg/d for 12 days (n = 4). Error bars represent SEM. *, $p < 0.05$; **, $p < 0.01$; ***, $p < 0.001$; ****, $p < 0.0001$ as analyzed by one-way ANOVA followed by post-hoc Holm-Šídák's test.

**Table S1. Information about the participants of this study, related to Figures 2 and S2.**

| # | DM† status | Sex | Age | Additional information | Fraction of OXT† neurons positive for $P2Y_{12}$ |
|---|---|---|---|---|---|
| I | No | Female | 69 | Malignant tumor of pancreas | No |
| II | No | Male | 59 | Colon cancer | No |
| III | Yes | Male | 77 | Upper gastrointestinal bleeding (brain subarachnoid hemorrhage), severe pneumonia, type 2 DM with nephropathy | 81.1% |
| IV | Yes | Female | 85 | DM with complications | 19.2% |
| V | Yes | Female | 74 | Gall bladder tumor, DM, 6 cardiac stents | 39.65% |

† DM, diabetes mellitus, OXT, oxytocin.

**Table S2. Information about male *M. fascicularis* used in this study, related to Figure 6.**

| # | Group | Age (y) | Initial weight (kg) | BCS[†] | Experiment |
|---|---|---|---|---|---|
| 1 | Cangrelor | 16 | 9.85 | 4.5 - 5 | 1 |
| 2 | Cangrelor | 16 | 10.85 | 4.5 - 5 | 1 |
| 3 | Cangrelor | 14 | 8.7 | 4 - 4.5 | 1 |
| 4 | Vehicle | 12 | 8.7 | 4 - 4.5 | 1 |
| 5 | Vehicle | 8 | 10 | 4.5 - 5 | 1 |
| 6 | Vehicle | 9 | 9.1 | 4.5 - 5 | 1 |
| 7 | Cangrelor | 12 | 8.66 | 4 - 4.5 | 2 |
| 8 | Cangrelor | 13 | 7.2 | 4 - 4.5 | 2 |
| 9 | Cangrelor | 10 | 7.7 | 4 - 4.5 | 2 |

[†] BCS, Body Condition Score.

**Table S3. Analysis of morning plasma parameters in male *M. fascicularis*, related to Figure 6.**

| Parameter | Units | Calorie restriction† | | | Cangrelor† | | |
|---|---|---|---|---|---|---|---|
| | | #7 | #8 | #9 | #7 | #8 | #9 |
| Total cholesterol | mmol/l | 3.7 | 3.25 | 2.11 | 2.94 | 1.45 | 3.1 |
| HDL | mmol/l | 1.81 | 1.06 | 1.39 | 1.65 | 1.1 | 1.85 |
| LDL | mmol/l | 1.86 | 1.78 | 0.97 | 1.29 | 0.76 | 1.66 |
| ApoA1 | g/l | 1.61 | 1.22 | 1.21 | 1.76 | 0.97 | 1.72 |
| ApoB | g/l | 0.38 | 0.34 | 0.2 | 0.38 | 0.36 | 0.71 |
| ApoA2 | mg/dl | 60.32 | 41.49 | 30.74 | 17.28 | 10.16 | 12.03 |
| ApoC2 | mg/dl | 1.49 | 1.2 | 0.76 | 0.99 | 1.03 | 0.85 |
| ApoC3 | mg/dl | 7.06 | 11.55 | 5.32 | 4.49 | 10.54 | 9.92 |
| ApoE | mg/l | 20.8 | 4.4 | 15.7 | 2.5 | 15.5 | 2.5 |
| LP-PLA2 | U/l | 78 | 78 | 62 | 51 | 48 | 145 |
| Homocysteine | µmol/l | 7.5 | 7.6 | 11.9 | 14.2 | 2.5 | 5.3 |
| Glucose | mmol/l | 4.81 | 4.97 | 4.34 | 3.78 | 4.05 | 4.2 |
| NT-proBNP | pg/ml | <10 | <10 | <10 | <10 | <10 | <10 |
| Phosphate | mmol/l | N/A | N/A | N/A | 1.53 | 1.12 | 1 |
| Calcium | mmol/l | N/A | N/A | N/A | 2.32 | 2.25 | 2.6 |
| Total bilirubin | µmol/l | N/A | N/A | N/A | 4.03 | 4.63 | 3.41 |
| Direct bilirubin | µmol/l | N/A | N/A | N/A | 1.24 | 1.27 | 0.4 |
| Total protein | g/l | N/A | N/A | N/A | 80.9 | 95.8 | 87.1 |
| ALT | U/l | N/A | N/A | N/A | 40.6 | 28.9 | 45.5 |
| AST | U/l | N/A | N/A | N/A | 32.4 | 44.1 | 33.5 |
| GGT | U/l | N/A | N/A | N/A | 73 | 43 | 40 |
| Cholinesterase | KU/l | N/A | N/A | N/A | 8.78 | 9.95 | 17.72 |
| Urea | mmol/l | N/A | N/A | N/A | 7.36 | 5.35 | 8.23 |

† Parameters in fasting morning plasma of animals #7, 8 and 9 before (last day before the termination of calorie restriction) and after receiving nasal cangrelor spray (Day 157, the last day before the termination of cangrelor administration). ALT, Alanine aminotransferase; Apo, apolipoproteins; AST, Aspartate aminotransferase; GGT, γ-Glutamyl transferase; HDL, high-density lipoprotein; LDL, low-density lipoprotein; LP-PLA2, lipoprotein-associated phospholipase A2; A2NT-proBNP, N-terminal pro-B-type natriuretic peptide.

**Movie S1, related to Figure 1.**

Representative speed up video recordings depicting ATP Inflares in the PVH regions-containing acute hypothalamic slices of male mice fed with chow or high-fat diet (HFD) for 2 weeks. Scale bar: 50 µm.

**Movie S2, related to Figure 1.**

Representative speed up video recordings depicting ATP Inflares in the PVH regions-containing acute hypothalamic slices of male mice fed with chow or high-fat diet (HFD) for 4 weeks. Scale bar: 50 µm.

**Movie S3, related to Figure 1.**

Representative speed up video recordings depicting ATP Inflares in the PVH regions-containing acute hypothalamic slices of male mice fed with high-fat diet (HFD) for 4 weeks before and after treatment with probenecid. Scale bar: 50 µm.

**Movie S4, related to Figure 1.**

Representative speed up video recordings depicting ATP Inflares in bilateral the PVH regions-containing acute hypothalamic slices of male *Panx1*$^{fl/fl}$ mice injected with AAV-GFAP-Cre (left side) or AAV-GFAP-mCherry (right side) and fed with high-fat diet (HFD) for 2 weeks. Scale bar: 50 µm.

**Movie S5, related to Figure 1.**

Representative speed up video recordings depicting ATP Inflares in the PVH regions-containing acute hypothalamic slices of male mice 2 wk after streptozotocin or vehicle injection. Scale bar: 50 µm.
